# Evaluation of CRISPR gene-editing tools in zebrafish

**DOI:** 10.1101/2020.10.19.345256

**Authors:** José M. Uribe-Salazar, Gulhan Kaya, Aadithya Sekar, KaeChandra Weyenberg, Cole Ingamells, Megan Y. Dennis

**Author notes:** Corresponding author: Megan Y. Dennis, Ph.D. University of California, Davis, School of Medicine, One Shields Avenue, Genome Center, 4303 GBSF, Davis, CA 95616.

## Abstract

**Background:** Zebrafish have practical features that make them a useful model for higher-throughput tests of gene function using CRISPR/Cas9 editing to create ‘knockout’ models. In particular, the use of G_0_ mosaic mutants has potential to increase throughput of functional studies significantly but may suffer from transient effects of introducing Cas9 via microinjection. Further, a large number of computational and empirical tools exist to design CRISPR assays but often produce varied predictions across methods leaving uncertainty in choosing an optimal approach for zebrafish studies.

**Methods:** To systematically assess accuracy of tool predictions of on- and off-target gene editing, we subjected zebrafish embryos to CRISPR/Cas9 with 50 different guide RNAs (gRNAs) targeting 14 genes. We also investigate potential confounders of G_0_-based CRISPR screens by screening control embryos for spurious mutations and altered gene expression.

**Results:** We compared our experimental *in vivo* editing efficiencies in mosaic G_0_ embryos with those predicted by eight commonly used gRNA design tools and found large discrepancies between methods. Assessing off-target mutations (predicted *in silico* and *in vitro*) found that the majority of tested loci had low *in vivo* frequencies (<1%). To characterize if commonly used ‘mock’ CRISPR controls (larvae injected with Cas9 enzyme or mRNA with no gRNA) exhibited spurious molecular features that might exacerbate studies of G_0_ mosaic CRISPR knockout fish, we generated an RNA-seq dataset of various control larvae at 5 days post fertilization. While we found no evidence of spontaneous somatic mutations of injected larvae, we did identify several hundred differentially-expressed genes with high variability between injection types. Network analyses of shared differentially-expressed genes in the ‘mock’ injected larvae implicated a number of key regulators of common metabolic pathways, and gene-ontology analysis revealed connections with response to wounding and cytoskeleton organization, highlighting a potentially lasting effect from the microinjection process that requires further investigation.

**Conclusion:** Overall, our results provide a valuable resource for the zebrafish community for the design and execution of CRISPR/Cas9 experiments.

## BACKGROUND

Zebrafish (*Danio rerio*) are increasingly used to rapidly and robustly characterize gene functions [1–4]. Features that make this model attractive over other classic vertebrate systems include external fertilization, rapid development, a large number of progeny, embryonic transparency, small size, and the availability of effective gene-editing tools [3, 5–12]. Continuous improvements of CRISPR editing in zebrafish have allowed efficient targeting of multiple genes simultaneously leading to rapid generation of either mosaic (G_0_) or stable mutant lines and subsequent characterizations of phenotypes [8, 13–19]. Such mutants have subsequently been used to test candidate genes associated with human diseases and developmental features [20]. The trend towards more affordable higher-throughput protocols using zebrafish requires a careful evaluation of methods used for the design of CRISPR-based genetic screens and potential confounders that may arise from the microinjection process that could artificially impact phenotypes.

New and creative CRISPR-based approaches in zebrafish address biological questions related to developmental processes (e.g., cell-lineage tracing) as well as gene functions (e.g., epigenome editing and targeted mutagenesis, reviewed in [21]). In the latter application, important factors in generating CRISPR gene knockouts include predicting/maximizing ‘on target’ Cas9 cleavage activity, predicting/minimizing unintended ‘off-target’ editing events, and rapidly detecting small insertions or deletions (indels). Presence of indels at candidate loci can be determined in an affordable manner via a number of approaches (reviewed in [22]), ranging from simple identification of heteroduplexes—arising from multiple alleles coexisting in the sampled DNA— visualized using a polyacrylamide gel electrophoresis (PAGE) [23] to more sophisticated sequencing approaches that precisely identify and quantify mutant alleles [14, 24]. On-target activity of a particular guide RNA (gRNA) can be predicted using tools that provide efficiency scores, often defined by information gathered across empirical assays [25]. One relevant example is CRISPRScan, a predictive-scoring system built from experimental zebrafish gene-editing data based on multiple factors such as nucleotide GC and AT content and nucleosome positioning [9, 26]. Bioinformatic tools also exist that define potential regions prone to off-target edits mainly based on sequence similarity and the type/amount of mismatches relative to the on-target region [26]. More recently, several methods have been devised to experimentally identify off-target cleavage sites (reviewed in [27]), including CIRCLE-Seq [28] and GUIDE-seq [29], that do not depend on prior sequence similarity information. These approaches are meant to provide a blind assessment of editing sites but do not necessarily reflect the *in vivo* activity of on-target activity of the CRISPR/Cas9 complex.

Previous studies have shown CRISPR off-target activity *in vivo* to be relatively low in zebrafish [8, 12, 18]. A cross-generational study identified no inflation of transmitted *de novo* single-nucleotide mutations due to CRISPR-editing using exome sequencing and a stringent bioinformatic pipeline [30] in a similar approach used to identify off-target mutations in mouse trios [31, 32]. Other studies have observed off-target mutation rates ranging from 0.07 to 3.17% in zebrafish by sequencing the top three to four predicted off-target regions based on sequence homology [11, 12, 18]. Although off-target mutations should not significantly impact studies of stable mutants, since unwanted mutations can be outcrossed out of studied lines relatively easily [14, 21], they may be problematic in rapid genetic screens using G_0_ mosaics that quickly test gene functions in a single generation.

The increasing number of tools available for the design and execution of CRISPR screens provide an important resource to the zebrafish community. Here, we assayed different available CRISPR on- and off-target prediction methods using empirical data from Cas9-edited zebrafish embryos. We quantified CRISPR cleavage efficiencies *in vivo* employing a variety of experimental approaches and used these results to compare the accuracy of *in silico* and *in vitro* tools for predicting Cas9 on- and/or off-target activity. Finally, to examine potential confounders that may arise from microinjection of Cas9 into embryos on resulting phenotypes, we assayed G_0_ ‘mock’ negative control embryos injected with a buffer containing either Cas9 enzyme or mRNA in the absence of gRNAs by performing RNA-seq and obtained a list of genes with significant differential expression versus uninjected wild-type siblings. In all, these results will serve as a useful resource to the research community as larger-scale G_0_ CRISPR screens become more common in assaying gene functions in zebrafish.

## RESULTS

### Identification of CRISPR-induced indels in zebrafish

We generated a dataset of experimentally confirmed indels within 14 protein-coding genes from injected NHGRI-1 wild-type zebrafish larvae targeted by 50 gRNAs (2–4 different gRNAs/target gene, assembled through the annealing of crRNA:tracrRNA) (Figure 1A, Supplementary Tables 1 and 2). These 50 gRNAs were designed using CRISPRScan [26] and include a range of predicted editing efficiencies (mean 57.6, range 23–83). To obtain experimental *in vivo* editing efficiency values for each gRNA, DNA extracted from a pool of 20 G_0_ mutant embryos — generated via microinjections of individual gRNAs at the one-cell stage and harvested at five days post-fertilization (dpf)— and ~200 bp regions surrounding predicted cut sites for all gRNAs were amplified and Illumina sequenced. To extract the proportion of reads carrying indel alleles, we used *CrispRVariants* [35] with uninjected batch siblings DNA as reference (Supplementary Table 2 for all scores obtained). From this, we inferred an *in vivo* ‘efficiency score’, calculated as the percentage of DNA from injected embryos harboring indels compared to uninjected batch siblings (Figure 1B).

**Figure 1.**
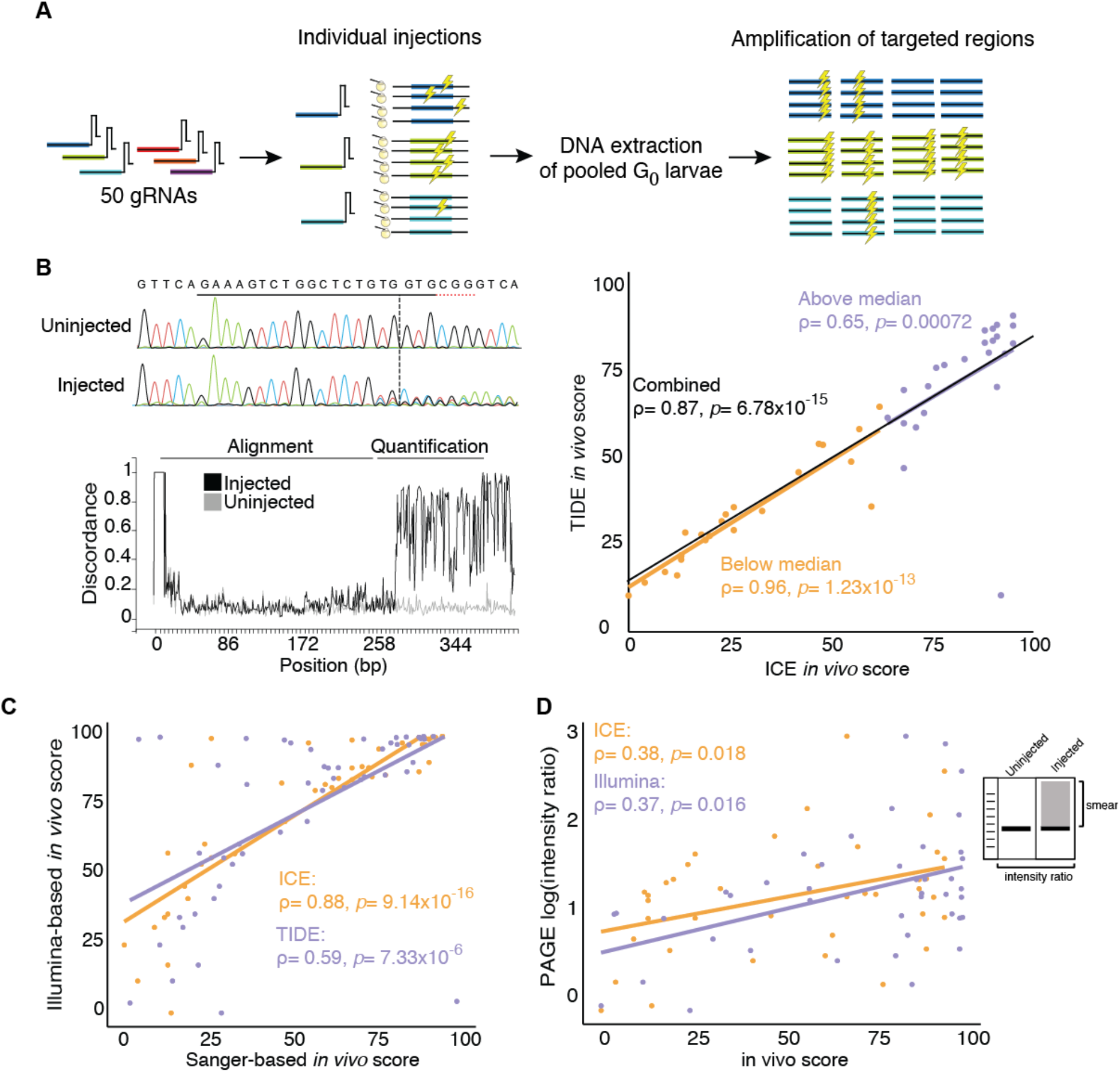
Workflow for the evaluation of CRISPR cleavages in NHGRI-1 zebrafish embryos. (**A)** The cartoon depicts our experiment, which included 50 gRNAs individually microinjected into one-cell stage embryos, DNA extracted from 20 pooled G_0_ larvae, and genomic regions targeted by the gRNA amplified. Lightning symbols represent a cleavage event. **(B)** An *in vivo* score was obtained from the Sanger sequencing traces using the ICE and TIDE tools, with an example output from ICE pictured. **(C)** Scores for the two tools were plotted with values below the median in orange and above the median in purple. **(D)** Scores from ICE and TIDE tools were compared to mosaicism percentages from Illumina sequencing of the same regions. **(E)** From the PAGE, an empirical intensity ratio was obtained and compared to the *in vivo* scores from Illumina and Sanger sequencing (ICE). Spearman correlations results are shown in the scatter plots with the line of best fit included.

To compare our efficiency scores with those produced from Sanger-based tools, we also amplified and sequenced ~500 bp fragments surrounding the targeted sites from the same DNA. We extracted the percentage of indels using two different tools that deconvolve major mutations and their frequencies within Sanger traces—Tracking of Indels by DEcomposition (TIDE) [33] and Inference from CRISPR Edits (ICE) [34] (Figure 1B). Briefly, these tools use the gRNA sequence to predict the cutting site in the control trace, map the sample trace to this reference, identify indels by deconvolving all base reads at each position, and provide a frequency of the indel spectrum [33, 34]. As previously reported [34], both tools provided positively correlated *in vivo* scores across all gRNAs (Spearman ρ= 0.87, *p*= 6.78×10^−15^) with an average score difference of 8.8±12.1 between tools (Figure 1C). We noted a higher correlation between tools in scores below the median (Spearman ρ= 0.96, *p*= 1.23×10^−13^) than above the median (Spearman ρ= 0.65, *p*= 0.00072; Figure 1C), suggesting that the deconvolution process in both tools is more accurate when fewer molecules from the pool carry indels. Both ICE and TIDE efficiency scores were correlated with our Illumina-based editing scores (ICE: Spearman ρ= 0.88, *p*= 9.14×10^−16^; TIDE: Spearman ρ= 0.59, *p*= 7.33×10^−6^; Figure 1D), though they significantly underestimated editing efficiencies with, for example, Illumina estimates 19.4±16.3 higher than ICE estimated scores (Figure 1D). Based on its higher correlation, we reported Sanger-based ICE *in vivo* scores for the rest of this study.

To ascertain consistency of editing efficiencies across embryos, we also repeated microinjections for four gRNAs targeting a single gene (*srgap2*) and assessed *in vivo* efficiencies scores of 20 individual larvae using the ICE tool. This resulted in low variance across injections and relative parity of efficiencies versus results from our pooled-larvae DNA preparations (e.g., low efficiency gRNA targeting exon 2; average±SE ICE score in individual larvae 18.9±3.3 versus an ICE score of 13 in pooled larvae and a high efficiency gRNA targeting exon 4; average±SE in individual larvae 69.2±5.3 ICE score versus an ICE score of 68 in pooled larvae).

A quicker and more affordable approach to quantify CRISPR cleavage efficiency is via PAGE, which takes advantage of the heteroduplexes produced from DNA harboring a mosaic mix of different types of indel mutations [23]. We performed PAGE on ~200 bp regions surrounding the predicted target site for each gRNA and quantified the PCR ‘smear’ intensity ratio of injected versus uninjected controls (see Methods). These intensity ratios were weakly correlated with our Illumina- (Spearman ρ= 0.37, p= 0.016) and ICE-estimated scores (Spearman ρ= 0.38, *p*= 0.018, Figure 1E), indicating that accurate quantitative efficiencies cannot be directly deduced from PAGE but that the intensity of PCR ‘smear’ does qualitatively convey CRISPR-cleavage efficiency.

### Accuracy of CRISPR on-target predictions by *in silico* methods

We next compared the accuracy of CRISPR on-target predictions computed by several published algorithms, including our chosen design tool CRISPRScan [26] (among the most popular tools used by the zebrafish community), CHOPCHOP [35–37] using two different scoring methods [38] [39] (among the most widely-used tool generally), E-CRISP [40], CRISP-GE [41], CCTop [42], CRISPRon [43], DeepSpCas9 [44], as well as the design tool from Integrated DNA Technologies (IDT, www.idtdna.com). Additionally, to assess if strain variability may have impacted our analysis—since all prediction tools used the Tübingen-derived reference genome (GRCz11) [4] whereas our study was performed in the NHGRI-1 strain (a cross between wild-type strains AB and Tübingen [46])—we obtained re-calculated CRISPRScan scores for our gRNAs using a modified zebrafish reference that included known NHGRI-1 variants [46] (now available as an additional reference in the tool browser at www.crisprscan.org). We then compared all *in silico* predicted efficiency scores to our *in vivo* mutagenesis from Illumina sequencing and ICE and observed a generalized underestimation of editing efficiency (Figure 2A). Strikingly, only scores predicted by CRISPRScan exhibited significant, albeit weak, correlation with *in vivo* scores. Further, the correlation with Illumina-based scores was significant only when using our NHGRIzed reference (Spearman ρ= 0.31, *p*= 0.028; Figure 2B), though CRISPRScan scores were highly concordant with those obtained using the Tübingen-derived reference (Spearman ρ= 0.88, *p*= 5.02×10^−17^; Figure 2B), with an average difference between scores of 4.2±4.6 (range 0–31) (Supplementary Table 2). CHOPCHOP values exhibited correlations with scores from four other *in silico* tools (E-CRISP, CRISPRon, DeepSpCas9, IDT) but none were concordant with our *in vivo* results (Figure 2B). Additionally, two tools that utilize deep learning methods (CRISPRon and DeepSpCas9) were significantly correlated with each other but failed to predict *in vivo* editing efficiencies in our assay (Figure 2B). Thus, despite the research community broadly adapting all methods for designing gRNAs, there is little consensus in predicting activity of a particular gRNA among these tools. Based on our results, we recommend using CRISPRScan for choosing gRNAs in zebrafish experiments.

**Figure 2.**
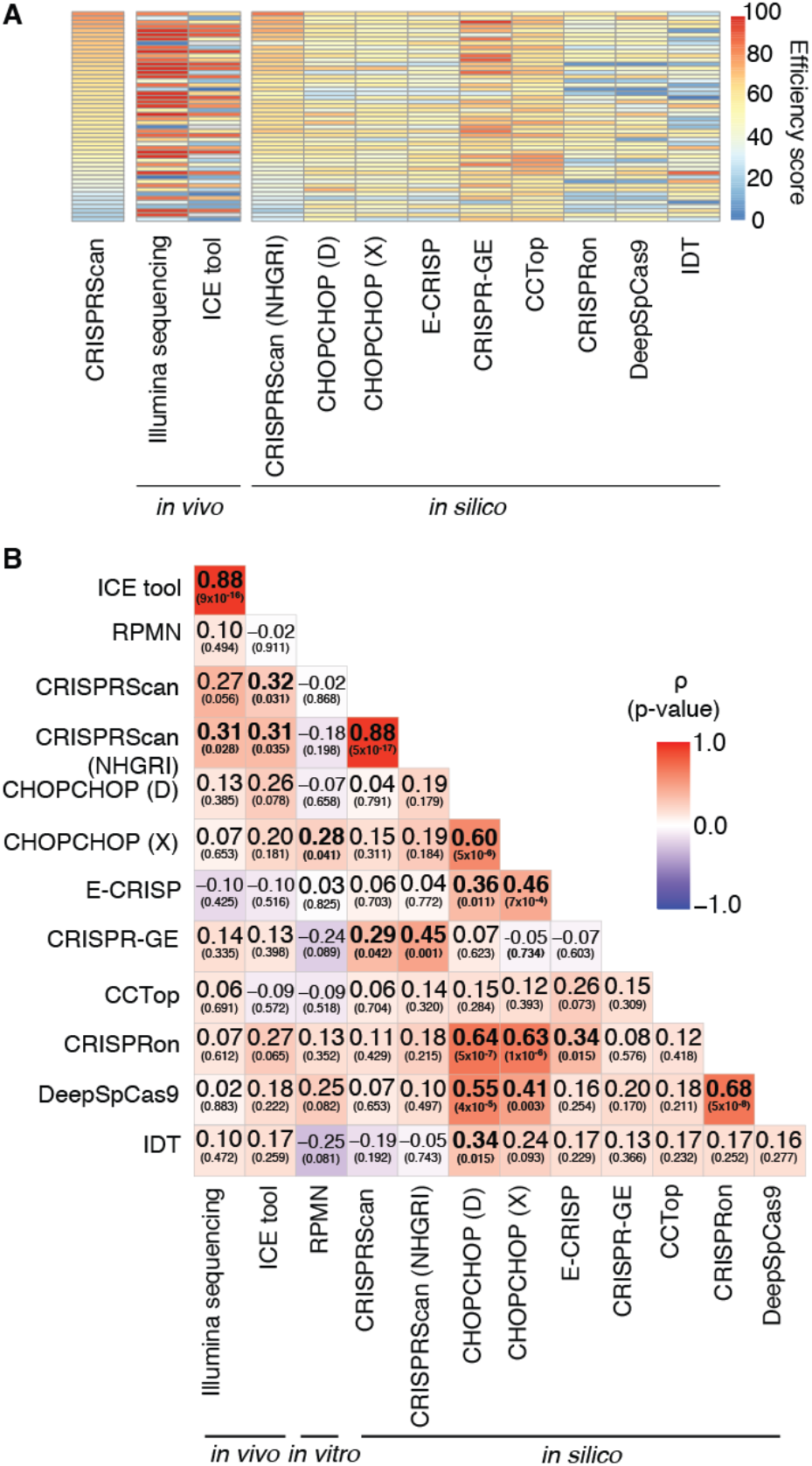
Correlation of on-target efficiencies calculated using different methods. **(A)** Heatmap of the efficiency scores obtained from the design tool (CRISPRScan), *in silico* prediction tools, and cutting cleavages obtained *in vivo* using Illumina sequencing and a deconvolution tool from Sanger sequencing [34] for 50 gRNAs. Each box represents a gRNA and the efficiency scores range from 0 (blue) to 100 (red). **(B)** Spearman correlations between all efficiency scores from *in silico* predictions, an *in vitro* protocol [28], and *in vivo* cutting assays. Each box includes the correlation result with the *p-value* in parenthesis. The color of the boxes represent the correlation values, ranging between −1 (blue) and 1 (red). CHOPCHOP scores were obtained using two different scoring methods, CHOPCHOP (D) (based on [39]) and CHOPCHOP (X) (based on [40]).

### Accuracy of CRISPR on-target predictions by an *in vitro* method

Next, we evaluated the possibility of using the *in vitro* protocol CIRCLE-seq [28], an approach designed to identify target sites of a given gRNA by subjecting naked genomic DNA to Cas9 enzyme/gRNA cleavage followed by Illumina sequencing, to obtain an editing efficiency score. It is important to emphasize that such *in vitro* assays are not designed for predicting on-target editing efficiencies. Nevertheless, we sought to understand if such an approach *could* be used for this purpose. We tested individually the 50 gRNAs described above using the CIRCLE-seq protocol [46], following the standard recommendations, and computed a log enrichment score normalized by the sequence library size, termed reads per million normalized (RPMN) (see Methods). We found that *in vitro*-obtained enrichment scores were not correlated with *in vivo* efficiencies (Illumina: Spearman ρ= 0.10, *p*= 0.494; ICE: Spearman ρ= −0.02, *p*= 0.911, Figure 2B) or with *in silico* predictions, with the exception of CHOPCHOP using the scoring method by [39] (Figure 2). This indicates that the CIRCLE-seq assay does not necessarily predict on-target CRISPR cleavage activity, at least quantitatively. Previous work from *in vivo* CRISPR studies of zebrafish suggests that increased GC-content predicts increased activity of gRNAs [26]. Examining GC content of our tested gRNAs, ranging from 31.8 to 77.3%, we observed a positive correlation with CRISPRScan *in silico* scores (linear model: beta= 68.18, *p*= 0.003, adjusted-r^2^= 0.16) and CIRCLE-seq *in vitro* RPMN scores (linear model: beta= 6.4, *p*= 0.006, adjusted-r^2^= 0.14) (Supplementary Figures 1A and B); however, our experimentally determined *in vivo* scores were not correlated with GC content (Illumina linear model: beta=−18.57, *p*= 0.689, adjusted-r^2^= −0.02; ICE linear model: beta=12.36, p= 0.817, adjusted-r^2^= −0.02; Supplementary Figure 1C), suggesting that additional variables should also be considered (e.g., depletion of A nucleotide bases, nucleosome positioning or DNA accessibility [26, 45]).

### CRISPR off-target mutation prediction methods

To avoid spurious phenotypes, off-target mutations should be minimized when choosing gRNAs in CRISPR experiments. To characterize off-target mutations for our set of 50 gRNAs, we queried predictions from *in silico* (CRISPRScan) and *in vitro* (CIRCLE-seq) methods. CRISPRScan provides a list of predicted off-target sites (between 55 and 1,350, median 206.5; Supplementary Table 3) for each gRNA within the zebrafish NHGRIzed reference genome (GRCz11/danRer11) based on a cutting frequency determination (CFD) score that primarily takes into account sequence similarity, location, and type of sequence mismatches [26, 38]. The CIRCLE-seq empirical approach also produced variable numbers of sites (between 18 and 874, median 113.5; Supplementary Table 3) per gRNA (defined as ‘CIRCLE-seq sites’) relative to the control library digested solely with Cas9 enzyme. The number of off-target sites predicted by CRISPRScan exhibited a significant, albeit weak, correlation with the number of CIRCLE-seq sites per gRNA (Spearman ρ=0.33, *p*= 0.022, Figure 3A). Focusing on putatively impactful off-target predictions, an average of 20±13% CRISPRScan-predicted and 64±7% CIRCLE-seq sites per gRNA intersected at least one gene (Supplementary Table 3). The sites predicted *in silico* or *in vitro* intersecting genes predominantly did not overlap with an average of 1.6±1.8 (range 0–7) genes per gRNA overlapping between the two approaches for the same gRNA.

**Figure 3.**
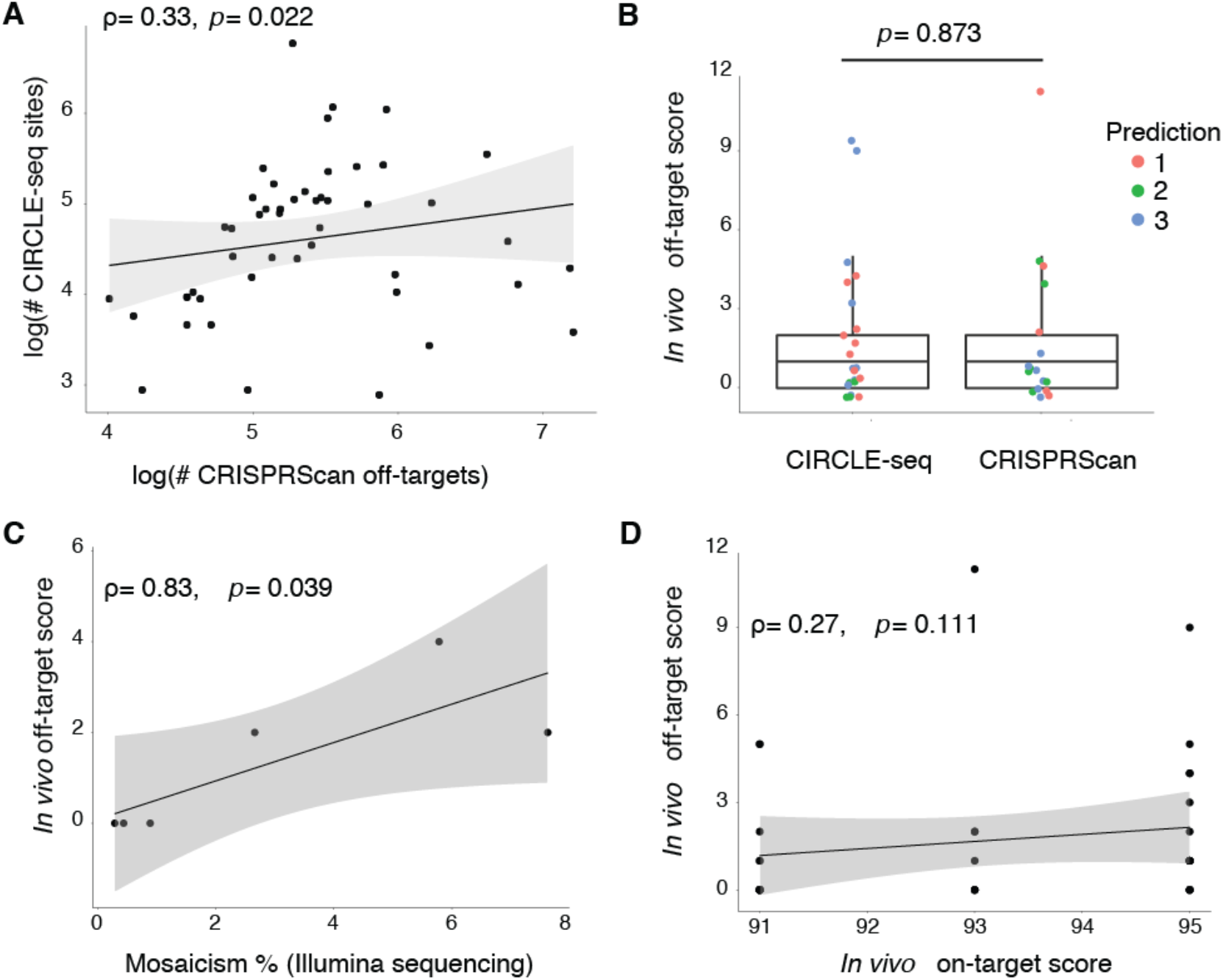
Assessment of off-target cleavage events using different prediction methods. **(A)** The number of predicted CRISPRScan off-target sites correlated with the number of identified CIRCLE-seq sites (Spearman correlation). Log normalization was used to reduce the range in the number of sites. **(B)** *In vivo* editing scores from the ICE tool for the top predicted off-target sites using CRISPRScan and CIRCLE-seq were not different. Scores were compared using a Mann-Whitney U test. **(C)** Editing efficiencies at predicted off-target sites using *in vivo* scores from Sanger sequencing and mosaicism % from Illumina sequencing were correlated (Spearman correlation). **(D)** Editing scores obtained *in vivo* at off-target sites were not correlated with the on-target efficiency of the gRNA. All scatter plots include the Spearman correlations results with the line of best fit.

To verify if predicted off-target sites were subjected to *in vivo* Cas9 cleavage, we performed Sanger sequencing of sites within genes identified *in silico* (n= 17) and *in vitro* (n= 20) for eight gRNAs, an average of six regions per gRNA (see Supplementary Table 1 for description of sites). Using the ICE tool, we found mosaic mutations at frequencies between 0 and 11%, with 23 out of the 37 sites evidencing indel frequencies below 1% (Figure 3B), and no differences observed between off-target sites predicted by CRISPRScan or CIRCLE-seq (Mann-Whitney U=175.5, *p*= 0.873; Figure 3B). To validate the accuracy of ICE at these low indel frequencies, we again performed Illumina sequencing of predicted off-target sites for six of the eight evaluated gRNAs (see Supplementary Table 1 for description) and found significant concordance in results (0.29–7.62% of mosaicism; Spearman ρ= 0.83, *p*= 0.039; Figure 3C). The average difference in mosaicism between ICE and Illumina was low (1.6±2.0), with ICE tending to slightly underestimate indel frequencies, highlighting its utility to quickly and economically assess predicted off-targets regions.

We also tested if sites predicted with higher likelihoods of off-target cutting events resulted in higher mutation rates by comparing the indel frequencies among the different levels of prediction (top 1, 2, or 3 prediction scores by CRISPRScan or CIRCLE-seq). No differences were found between prediction groups (Kruskal-Wallis: H_(2)_= 2.26, *p*= 0.320; Figure 3B), suggesting that the information used by the tools to assign probabilities of off-target activity (e.g., CFD scores in CRISPRScan or normalized read counts in CIRCLE-seq) do not necessarily predict the efficiency of cutting at off-target sites *in vivo*. Thus, off-target cutting mutations at the assessed sites exhibited low frequencies with no clear method performing best. Moreover, none of the on-target scores previously obtained (*in silico*, *in vitro*, or *in vivo*) correlated with the number of *predicted* off-target sites per gRNA (using either CRISPRScan or CIRCLE-seq), nor the frequency of indels at validated off-target sites (Spearman ρ= 0.27, *p*= 0.111, Figure 3D), suggesting that higher on-target efficiencies do not necessarily translate into increased frequencies of spurious off-target mutations.

### Evaluating CRISPR Cas9-injection controls

A commonly used ‘mock’ injection control for phenotypic screens of CRISPR-generated G_0_ mosaic lines are embryos injected with buffer and Cas9 in the absence of a gRNA. We sought to determine if such control treatments could significantly impact the genome or transcriptome of our zebrafish larvae. To characterize its impact on genes, we performed RNA-seq of wild-type NHGRI-1 embryos injected with either Cas9 enzyme or Cas9 mRNA (three biological replicates of a pool of five injected larvae), uninjected batch siblings (two biological replicates of a pool of five larvae), and uninjected siblings from another batch (three biological replicates with a pool of five larvae) as controls.

#### Potential genomic mutations in controls

Recently, Sundaresan and colleagues [46] found that Cas9 in the presence of Mn^+2^ ions can result in double-strand cleavage of genomic DNA in the absence of a gRNA. Although their study did not show this same off-target cleavage activity in the presence of Mg^+2^, we hypothesized that aberrant genomic mutations could be incurred by Cas9 due to the presence of MgCl_2_ in our injection buffer since Mg^+2^ has been shown to compete with Mn^+2^ in activating common enzymes [47]. Using our RNA-seq data, we used an optimized pipeline [48] to identify somatic mosaic mutations with uninjected wild-type controls as a reference for common polymorphisms. Focusing only on high-confidence variants (minimum sequence read depth of 20), we filtered already-reported variants in the NHGRI-1 zebrafish line [49], and used the Variant Effect Predictor tool from ENSEMBL to obtain a list of frameshift mutations in protein-coding genes present in our Cas9-injected larvae. A total of 48 and 38 genes were identified with frameshifting variants in larvae injected with Cas9-enzyme and Cas9-mRNA, respectively, with 14 of these genes shared across both injection types (Figure 4A, Supplementary Table 4). On average, each pool of larvae injected with Cas9 enzyme or mRNA carried frameshift variants in 18.7±3.1 genes. All identified frameshift variants evidenced low allelic frequencies (Cas9-enzyme: average 0.043, range 0.0036-0.142; Cas9-mRNA: average 0.059, range 0.002-0.316) and high read depth (Cas9-enzyme: average 386.5, range 22-2076; Cas9-mRNA: average 343.8, range 20-3453) (Supplementary Table 4). Additionally, frameshift variants were positioned closer to a potential Cas9 PAM site (NGG) than by random chance (4 bp median observed distance to closest PAM site; empirical *p*= 0.0016 using the whole-genome and *p*= 0.006 using protein-coding regions only, from 10,000 permutations). Therefore, we decided to evaluate if indels would consistently arise in these genes in an additional set of microinjections.

**Figure 4.**
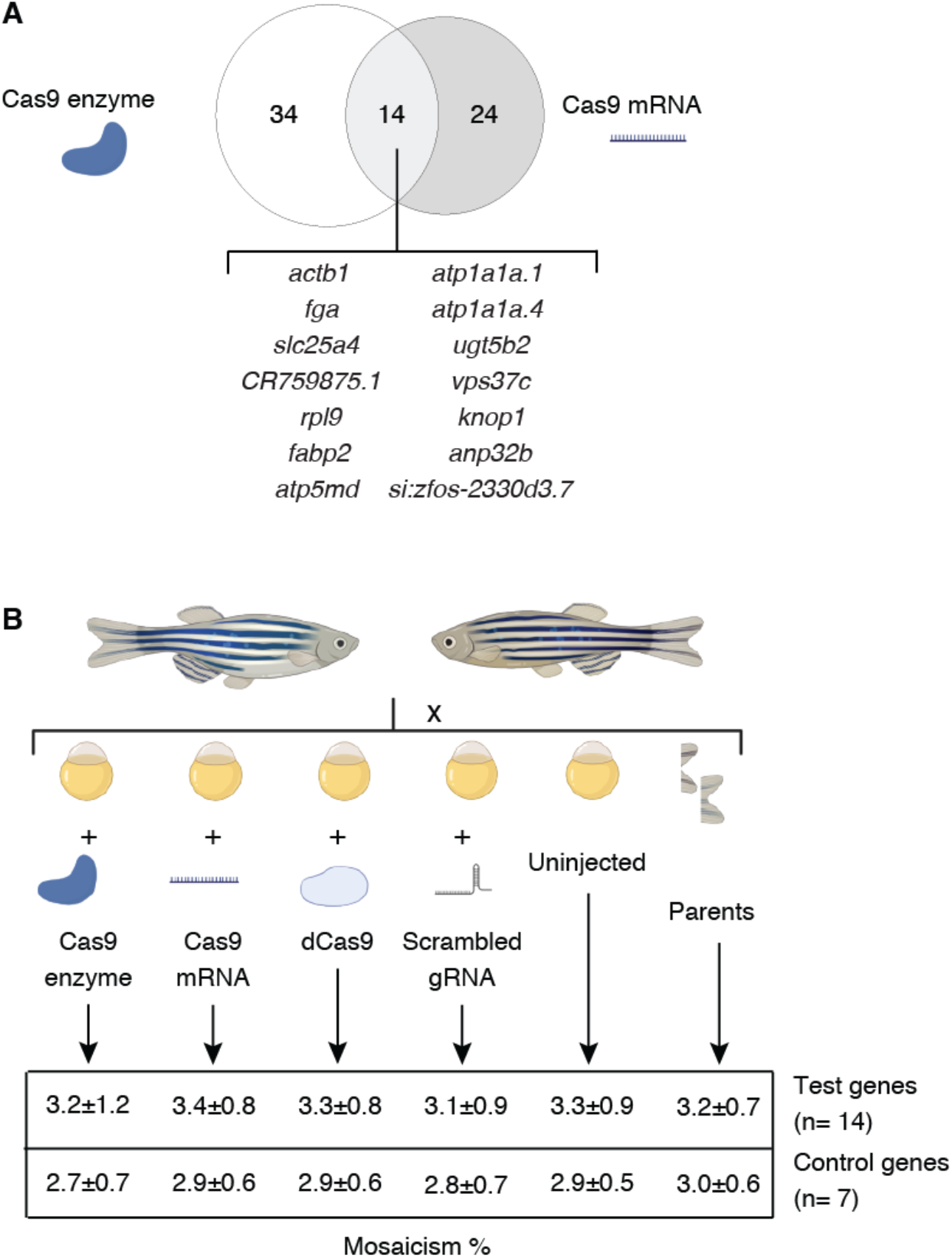
Evaluation of spurious genomic mutations in CRISPR-injection controls. **(A)** The abundance of protein-coding genes carrying frameshift variants for each Cas9-injected treatment are depicted in a Venn diagram, with mutated genes identified in both treatments listed. **(B)** Genomic DNA from zebrafish larvae injected with Cas9 enzyme, Cas9 mRNA, catalytically dead Cas9 (dCas9), a scrambled gRNA, uninjected batch siblings, and a fin clip from their parents was used to perform targeted Illumina sequencing of 21 genes to quantify indel mosaicism with average ± standard deviation values listed in the table (see Supplementary Table 1 and Supplementary Table 7 for the description of the genes).

We performed a new set of microinjections in NHGRI-1 larvae using these same controls (Cas9 enzyme and Cas9 mRNA) and two additional ones commonly used in CRISPR experiments (catalytically dead Cas9 (dCas9) enzyme and a scrambled gRNA coupled with Cas9 enzyme, sequence published in [19]) and evaluated the presence of mutations in 21 genes, including 14 genes with identified frameshift mutations in our RNA-seq data and seven controls with no mutations observed (Figure 4B, see Supplementary Table 1 and Supplementary Table 5 for the description of all sites). Briefly, genomic DNA was harvested from (1) three pools of five larvae from each group injected at the one-cell stage (Cas9 enzyme, Cas9 mRNA, dCas9, scrambled gRNA); (2) three pools of five uninjected batch siblings larvae; and (3) finclips of the crossing parents as controls. Subsequently, ~200 bp regions surrounding the closest Cas9 PAM site to the previously RNA-seq-identified variants were Illumina sequenced and the alleles extracted using *CrispRVariants* [50]. We did not observe evidence of inflation of indels in any of the injected groups relative to the uninjected batch siblings or the parental fish, with an overall average mosaicism of 3.1±0.8% per site (below the expected 10% allele ratio for a heterozygous variant in a single individual from a pool of five; Figure 4B, Supplementary Table 5). Our NHGRI-1 zebrafish carried common single nucleotide variants in the targeted regions, particularly in gene *si:ch1073-110a20* where two variants were present in close to 50% and 20% of the reads (Supplementary Figure 2). Interestingly, we did observe a subtly higher mosaicism in the genes previously detected with variants in our RNA-seq data relative to the regions used as controls (Mann-Whitney U= 2251.5, *p*= 0.00074, median mosaicism in tested genes 3.4%, median mosaicism in control genes 2.88%; Figure 4B, Supplementary Table 5). Thus, it is possible that the genes we identified with variants in our RNA-seq data may be naturally prone to carry variants. In summary, these results suggest that currently used CRISPR controls do not suffer systematic DNA cleavages in the absence of a gRNA.

#### Differential gene expression in controls

We also characterized the impact of injecting Cas9 enzyme or mRNA on the transcriptomes of our zebrafish larvae. Comparisons of transcripts abundances show significant variance across biological replicates when quantifying in both Cas9 treatments, particularly evident in samples injected with the Cas9 enzyme, versus wild-type uninjected larvae (Figure 5A). This suggests that considerable stochasticity may exist regarding the effects of Cas9 injections in these controls. Examining the genes impacted, we identified hundreds of differentially-expressed (DE) genes in our Cas9-injected versus uninjected controls, with a greater number of upregulated genes than downregulated genes (Figure 5B, Supplementary Table 6). Specifically, Cas9-enzyme injections resulted in a total of 1,100 DE genes (3.6% of the genes assayed), with 756 genes (68.7%) upregulated (fold change > 1) and 344 (31.3%) downregulated (fold change < −1). Cas9-mRNA injected larvae exhibited 548 DE genes (1.8% of the genes assayed), 376 (68.6%) of these upregulated and 172 (31.4%) downregulated (Figure 5B). We observed 248 (197 upregulated and 51 downregulated) common DE genes between the two treatments (Figure 5C), which could be part of a common response to the microinjection process. Network analyses identified commonalities in the shared DE genes enriched in key regulators of different KEGG pathways, including spliceosome and ribosome (including genes *eif4g2b*, *eif4g1a*, *hnrnpd*, *magoh*, *hnrnpa0a*), hedgehog signaling (*shha*), glutathione metabolism (*gsto2*, *gsr*), GnRH signaling (*dusp6*), aminoacyl-tRNA biosynthesis (*yars*), cell cycle (*kif2c*), glycolysis (*aldoca*), and cellular senescence (*ppp3cca*) (Supplementary Figure 3). Furthermore, while we observed no enrichment in gene ontology terms for downregulated genes, common upregulated genes from both treatments were related to response to wounding (GO:0009611, adjusted *p*-value= 0.009) and cytoskeleton organization (GO:0045104, adjusted *p*-value=0.009) (Supplementary Table 7), revealing molecular consequences of the microinjection process that were still detectable five days later.

**Figure 5.**
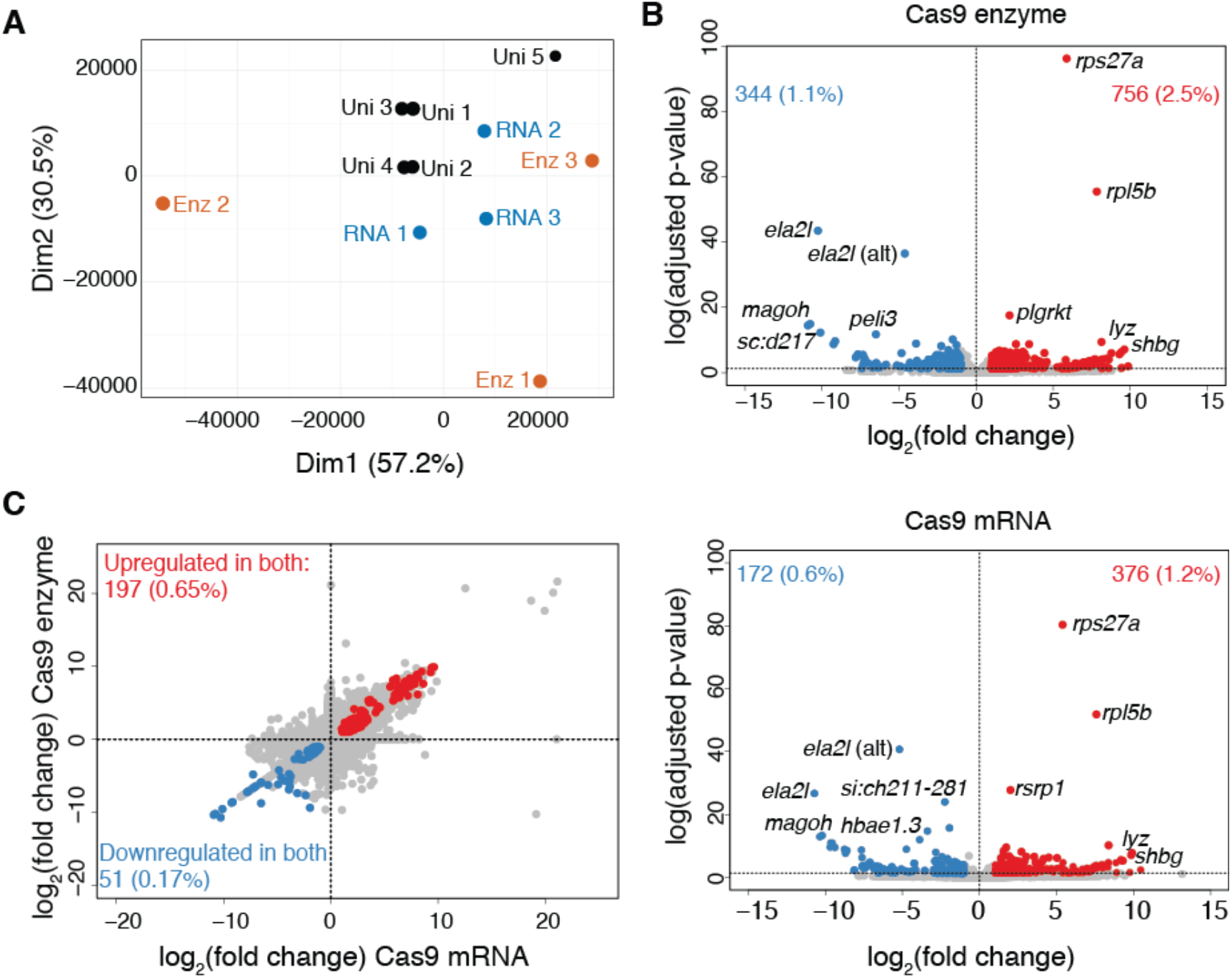
Evaluation of expression variability in CRISPR-injection controls. **(A)** Principal components analysis using the transcript abundances in larvae injected with Cas9 enzyme (Enz1, Enz2, Enz3), Cas9 mRNA (RNA1, RNA2, RNA3), uninjected siblings (Uni1, Uni2), and uninjected siblings from a different batch (Uni3, Uni4, Uni5). **(B)** Volcano plots show the differentially-expressed genes in Cas9-enzyme and Cas9-mRNA injected larvae with the number (and %) of upregulated (fold change > 1) and downregulated (fold change < −1) genes. The top five representative up- and downregulated genes are highlighted, with the full list of genes available as Supplementary Table 6. **(C)** Differentially-expressed genes across samples injected with Cas9 enzyme or Cas9 mRNA relative to uninjected batch-siblings show significant correlations. Plots include the numbers and percentages (in parentheses) of genes downregulated (blue) and upregulated (red) in both Cas9 treatments from the total amount of genes assayed (n= 30,258).

## DISCUSSION

Our study presents a comprehensive evaluation of empirical and predictive tools currently used for CRISPR editing in zebrafish. Cleavage scores obtained by an *in vivo* assessment of 50 gRNAs via Sanger sequencing and deconvolution tools (ICE and TIDE) were concordant with Illumina sequencing, the gold standard in predicting efficiencies, as previously reported [34]. Both tools underestimated the presence of non-edited alleles by ~20%, contrary to previous comparisons of TIDE and Illumina sequencing in cell lines, where TIDE showed a ~10–20% overestimation of non-edited alleles [51]. For sites with lower indel frequencies, as we observed for predicted off-target mutations, ICE scores were more concordant with Illumina results (~1– 2% difference, again mostly underestimates). Therefore, we suggest that Sanger sequencing deconvolution tools are valuable for establishing relative gRNAs efficiencies but do not necessarily accurately predict absolute cleavage efficiencies in zebrafish *in vivo*, except at sites with low indel frequencies. In addition, we formalized an empirical ‘intensity ratio’ score from the commonly-used PAGE approach to assay CRISPR indels and verified its utility in approximating cleavage efficiencies, making it a more affordable and rapid approach to assay editing efficiencies versus sequencing.

On-target efficiency prediction tools showed large differences using the same set of gRNAs sequences, highlighting the importance of understanding features accounted for by each tool. A recent review [25] provides a comprehensive overview of different design tools available and the source of experimental data used to train each one. CRISPRScan [26] was the only tool that could predict on-target efficiency in our set of gRNAs, while no other method provided scores that were correlated with cleavage activities observed *in vivo*. One limitation of our study was the skew in higher efficiency gRNAs (mean predicted CRISPRScan score of 57.6), which could feasibly impact correlations. Notably, we did obtain more accurate CRISPRScan predictions when we utilized our NHGRIzed reference [49] compared to the current Tübingen-derived reference [4], highlighting the importance of accounting for known genetic variation when designing suitable gRNAs [52, 53]. Considering CRISPRScan was the only tool that incorporated empirical data from zebrafish, with most methods tested using *in vitro*-derived data, our results emphasize the importance of utilizing a tool trained using *in vivo* experimental data specific to the study’s target species.

An *in silico* (CRISPRScan) and *in vitro* (CIRCLE-seq) method predicted ~20% and 65% potential off-target regions impacting genes, respectively. Notably, we did not evaluate if other predicted sites included *cis*-regulatory elements that could also potentially alter gene expression. Future assessments should include tests targeting a diversity of loci for a more thorough understanding of the potential off-target indels caused by unwanted CRISPR cleavage sites. We observed low off-target mutation frequencies (most <1%), similar to those previously reported from using single [11, 12] or multiple gRNAs [18], although did observe off-target indel frequencies as high as 11% for certain gRNAs. Notably, neither predictive method (CRISPRScan or CIRCLE-seq) nor their likelihood score (using CFD or normalized read count) could accurately predict indel frequencies at off-target sites. Typically, such low mutation frequencies should not be of high impact when generating stable knockout zebrafish lines as these could be easily outcrossed. However, such mutations could have significant impacts on phenotypic outcomes when injected G_0_ mosaic populations are analyzed directly.

The adequate selection of controls is a fundamental process in evaluating gene function using G_0_ knockout crispant zebrafish, as these larvae serve as baselines from which inferences will be made from. Currently, no consensus exists for preferred controls used in high-throughput CRISPR workflows of zebrafish larvae, which can include targeting a known gene as a positive control (e.g., *tyr*) [14], uninjected larvae [17, 18], sham injections with a Cas9:tracrRNA complex [15], and injections of a scrambled gRNA [16, 19], among others. Our RNA-seq assay identified several genes carrying frameshift mutations using uninjected clutch siblings as reference. A follow-up analysis of a second set of injections showed existence of mosaic variants in all injected controls (e.g., Cas9 mRNA, enzyme, and scrambled gRNA), in addition to uninjected siblings and crossed parents at low allelic frequencies (~3%). Nevertheless, even though we were limited to our targeted regions, we did observe a higher mosaicism in genes identified as carrying frameshift mutations from our RNA-seq assay compared to control genes, suggesting that these genes could be naturally prone to exhibit mutations in the NHGRI-1 zebrafish line. We also observed high variability in gene expression in larvae solely injected with Cas9 enzyme or mRNA, with several of these DE genes involved in response to wounding processes. Notably, these DE genes were retrieved from 5 dpf larvae suggesting that damage incurred during the microinjection process has a lasting effect. These results suggest that caution should be taken in using G_0_ mosaic mutants in investigating phenotypes related to pathways found to be significantly skewed in injection controls, including those involving the function of the spliceosome, ribosomes, and cytoskeleton dynamics.

## CONCLUSIONS

Overall, we performed a simultaneous assessment of gRNA activities predicted by several commonly used *in silico* and *in vitro* methods with those determined experimentally *in vivo* in injected zebrafish embryos. These results provide valuable information that can be incorporated into the design and execution of CRISPR/Cas9 assays in zebrafish using available workflows [8, 13, 14, 17, 18]. Namely, we make the following conclusions and recommendations:

- Sanger-based efficiency estimates (TIDE and ICE) tend to underestimate indel mosaicism in zebrafish, though they are more accurate when lower mutational mosaicism exists (such as those observed at off-target sites).
- Quantifying heterodimers via PAGE gels represents an affordable method to qualitatively assay CRISPR cutting efficiencies.
- Of the existing tools, we recommend CRISPRScan for predicting gRNA on-target efficiency, preferably matched to the zebrafish strain being used.
- Off-target mutations occur at relatively low rates with neither *in silico* nor *in vitro* prediction methods performing significantly better.
- Microinjection of Cas9 (enzymes or mRNA) into embryos does not result in spurious genomic mutations but does impact certain genes and pathways. Caution should be exercised if studying phenotypes related to these genes when performing G_0_ mosaic zebrafish screens.

Our aim was to provide information to aid in the decision-making process for future projects using affordable and reliable gene-editing tools in zebrafish. As higher-throughput methods continue to be developed for assaying multiple genes simultaneously, it will be important to use optimal tools for predicting and assessing on- and off-target activity in zebrafish larvae for accurate interpretation of phenotypic outcomes.

## METHODS

### Zebrafish husbandry

NHGRI-1 wild-type zebrafish [49] were maintained through standard protocols [54] and their use was approved by the Institutional Animal Care and Use Committee from the Office of Animal Welfare Assurance, University of California, Davis. Animals were kept in a temperature (28±0.5°C) and light (10 h dark/14 h light cycle) controlled modular system with UV-sterilized filtered water (Aquaneering, San Diego, CA), with a density of 25 adult fish per tank. Feeding and general monitoring of all zebrafish was performed twice a day (9 am and 4 pm). Food included rotifers (Rotigrow Nanno, Reed Mariculture, Campbell, CA), brine shrimp (Artemia Brine Shrimp 90% hatch, Aquaneering, San Diego, CA), and flakes (Zebrafish Select Diet, Aquaneering, San Diego, CA). For all experimental procedures, eggs were collected via natural spawning of randomly selected adult NHGRI-1 zebrafish in 1 liter crossing tanks (Aquaneering, San Diego, CA), using a minimum of five breeding pairs (1 male, 1 female) unless otherwise specified. Embryos were grown in standard Petri dishes with E3 media (0.03% Instant Ocean salt in deionized water) and incubated at 28±0.5°C, using a dissecting microscope (Leica, Buffalo Grove, IL) for developmental staging and daily monitoring until their use for molecular procedures.

### Design and *in silico* predictions for gRNAs

50 gRNAs targeting exons of 14 genes were designed using CRISPRScan [26] (scores ranging between 24 and 83 with a mean value of 57.6) with zebrafish genome version GRCz11/danRer11 as the reference (see description of gRNAs in Supplementary Tables 1 and 2). All targeted genes were protein coding. For each designed gRNA, we obtained the efficiency scores predicted by CRISPRScan [26], CHOPCHOP [35–37] using the scoring method from [38] and [39], E-CRISP [40], CRISPR-GE [41], CCTop [42], CRISPRon [43], DeepSpCas9 [44], and the IDT design tool (www.idtdna.com). From CRISPRScan, we also gathered the predicted off-target sites for each gRNA defined by the CFD score [26]. Additionally, we utilized *bedtools* [55] to determine the GC percentage for each gRNA. To incorporate NHGRI-1 variants into the zebrafish reference, we used the FastaAlternateReferenceMaker function from *GATK* [56] with the reported high-confidence variants for the NHGRI-1 zebrafish strain [49].

### Microinjections to generate CRISPR G_0_ mosaic mutants

All gRNAs were individually injected into NHGRI-1 embryos to estimate the frequency of indels. gRNAs were prepared following the manufacturer’s protocol (Integrated DNA Technologies). Briefly, 2.5 μl of 100 μM crRNA, 2.5 μl of 100 μM tracrRNA, and 5 μl of Nuclease-free Duplex Buffer using an annealing program consisting of 5 min at 95°C, a ramp from 95°C to 50°C with a −0.1°C/s change, 10 minutes (min) at 50°C, and a ramp from 50°C to 4°C with a −1°C/s change. Ribonucleoprotein injection mix was prepared with 1.30 μl of Cas9 enzyme (20 μM, New England BioLabs), 1.60 μl of prepared gRNAs, 2.5 μl of 4x Injection Buffer (containing 0.2% phenol red, 800 mM KCl, 4 mM MgCl_2_, 4 mM TCEP, 120 mM HEPES, pH 7.0), and 4.6 μl of Nuclease-free water. Microinjections directly into the yolk of NHGRI-1 embryos at the one-cell stage were performed as described previously [57], using needles from a micropipette puller (Model P-97, Sutter Instruments) and an air injector (Pneumatic MPPI-2 Pressure Injector). Embryos were collected and ~1 nl of ribonucleoprotein mix was injected per embryo, after previous calibration with a microruler. Twenty injected embryos per Petri dish were grown up to 5 dpf at 28°C.

### Illumina and Sanger amplicon sequencing

DNA extractions were performed on 20 pooled embryos by adding 100 μl of 50 mM NaOH, incubation at 95°C for 20 min, ramp from 95°C to 4°C at a 0.7°C/s decrease, followed by an addition of 10 μl of 1 M Tris-HCl and a 15 min spin at 4680 rpm. We amplified a ~200 bp region surrounding the targeted site of each gRNA (see Supplementary Table 1 for description of primers). PCR amplifications were performed using 12.5 μl of 2X DreamTaq Green PCR Master Mix (Thermo Fisher), 9.5 μl of Nuclease-Free water, 1 μl of 10 μM primers, and 1 μl extracted DNA. Thermocycler program included 3 min at 95°C, followed by 35 cycles of 15 s at 95°C, 30 s at 60°C, and 20 s at 72°C, and a final 5 min incubation at 72°C. Reactions were purified using Ampure XP magnetic beads (Beckman Coulter) and Illumina sequenced (Genewiz, San Diego, CA). To obtain percent mosaicism of mutants by mapping paired-end fastq reads to the zebrafish reference genome (GRCz11/danRer11) using *bwa* [57] and the R package *CrispRVariants* [35]. Additionally, we amplified a ~500 bp region surrounding the targeted site of each gRNA from the same extracted DNA for six gRNAs and performed and performed Sanger sequencing (Genewiz, San Diego, CA). Raw trace files were used in the TIDE [33] and ICE [34] tools to predict the percentage of indels, which we used as our *in vivo* editing score for each gRNA. For both Sanger and Illumina sequencing, we used uninjected batch-sibling embryos as a control reference.

### PAGE and intensity-ratio estimation

An empirical cleavage analysis from each gRNA was performed using PAGE. Briefly, we amplified a ~200 bp region in DNA around the targeted site from gRNA-injected and uninjected embryos, as described above. Reactions of the uninjected and injected samples from the same amplicon were run on a 7.5% polyacrylamide gel together for 75 min at 110 V and revealed using GelRed (VWR International). Gel images were processed in the software Fiji [58]. For each sample, we defined areas **A** and **B** as follows:

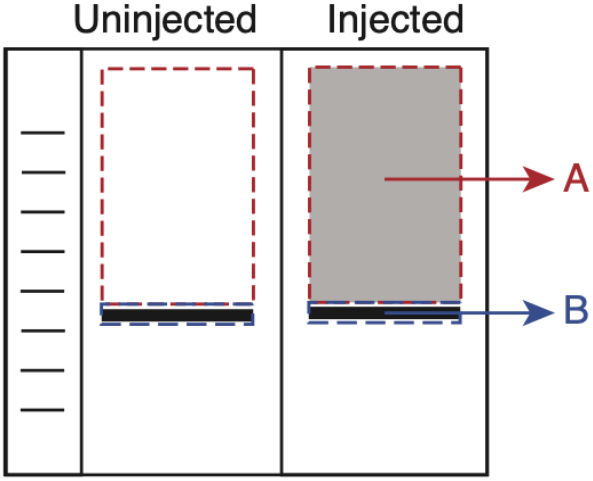

For each gRNA, the mean-intensity value was obtained for the A and B areas in both the injected and uninjected samples. The A and B areas were exactly the same size between samples. The intensity ratio was calculated as: [injected B / injected A] / [uninjected B / uninjected A]. Log-normalized intensity ratios followed a normal distribution (Shapiro-Wilk test: W= 0.96, *p*= 0.167) with an average value of 1.21±0.70.

### CIRCLE-seq

CIRCLE-seq libraries were prepared for each gRNA (IDT) using genomic DNA extracted from NHGRI-1 (DNA Blood & Tissue kit, Qiagen) following the described protocol [59]. Libraries were sequenced using one HiSeq XTen lane (Novogene, Sacramento, CA), providing an average of 7.3 million reads (range: 4.0 - 13.3 million reads) and >Q30 for 92% of reads per gRNA library. Raw reads were processed using the bioinformatic pipeline described [59] (mapping rate >99% in all samples) to identify regions with cutting events relative to a control sample (treated with Cas9 enzyme and no gRNA). In an attempt to obtain an on-target efficiency estimation from *in vitro* digestions, we calculated the reads per million normalized (RPMN). For this purpose, we used *samtools* [60] to extract read coverage from aligned bam files. For each gRNA, coverage was obtained for the third and fourth base upstream of the PAM site as it is the region expected to be cut by Cas9 [61]. RPMN for each gRNA was calculated as the sum of coverage at these two sites divided by the total mapped reads per sample and multiplied by one million to scale the values. RPMN scores ranged from 4.42 to 881 (median 99.3) so we decided to use a log normalization to reduce this range.

### RNA-seq

We performed RNA-seq of Cas9 injected NHGRI-1 larvae to identify potential gRNA-independent cleavage sites. One-cell stage NHGRI-1 embryos were injected with either Cas9 enzyme or Cas9 mRNA. Injection mix for Cas9 enzyme included Cas9 enzyme (20 μM, New England BioLabs), 2.5 μl of 4x Injection Buffer (0.2% phenol red, 800 mM KCl, 4 mM MgCl_2_, 4 mM TCEP, 120 mM HEPES, pH 7.0), and Nuclease-free water. Cas9 mRNA was obtained from plasmid pT3TS-nCas9n (Addgene, plasmid #46757) [5], using the MEGAshotscript T3 transcription kit (Thermo Fisher) following manufacturer’s guidelines of 3.5 h 56°C incubation with T3. mRNA was purified with the MEGAclear transcription clean-up kit (Thermo Fisher) and concentration of mRNA obtained using a NanoDrop (Thermo Fisher). The injection mix of Cas9 mRNA contained 100 ng/μl of mRNA, 4x Injection Buffer (0.2% phenol red, 800 mM KCl, 4 mM MgCl_2_, 4 mM TCEP, 120 mM HEPES, pH 7.0), and Nuclease-free water.

Additionally, uninjected batch-siblings and uninjected siblings from an additional batch were used as controls. All embryos were grown at 28°C in a density of <50 embryos per dish. At 5 dpf, three pools of five larvae were collected for each group (Cas9 enzyme, Cas9 mRNA, and uninjected) for RNA extraction using the RNeasy kit (Qiagen) with genomic DNA eliminator columns for DNA removal. Whole RNA samples were subjected to RNA-seq using the poly-A selection method (Genewiz, San Diego, CA).

### Variant identification from RNA-seq data

We followed a previously described pipeline to identify somatic variants from RNA-seq data [48]. Briefly, we mapped reads with *STAR* [62] using the 2-pass mode and a genomic reference created with GRCz11/danRer11 assembly and gtf files (release version 100). Variant calling was performed with MuTect2 as part of *GATK* [56] using the tumor versus normal mode. ‘Normal’ was defined by the two uninjected samples to identify all somatic mutations in our Cas9 injected embryos. Variants were annotated using the Variant Effect Predictor tool [63]. High confidence variants (minimum sequencing depth of 20) previously reported for the NHGRI-1 line [49] were removed. Only frameshift loss-of-function variants with a minimum read depth of 20 in canonical protein-coding genes were considered. We extracted the median distance between the identified variants and the nearest Cas9 PAM site (NGG sequence) using the coordinates in the CRISPRScan UCSC track. This median observed distance was compared to the result of median distances of 10,000 permutations of random sampling across the genome and their nearest PAM site. One-tailed empirical *p* values from this comparison were calculated as (M+N)/(N+1), where M is the number of iterations with a median distance below the observed value and N is the total number of iterations. We orthogonally investigated the presence of variants in 23 genes via Illumina sequencing of a ~200 bp region surrounding the identified variant location and the R package *CrispRVariants* [50] (Supplementary Table 1 for primers description). For this purpose, we extracted DNA from 3 pools of 5 embryos injected with Cas9 enzyme, Cas9 mRNA, dCas9 (Alt-R S. p. dCas9 protein V3 from IDT), a scrambled gRNA (see Supplementary Table 1 for sequence description), or uninjected. In addition, we extracted DNA from a finclip of the crossing parents of the embryos used for the injections (both female and male). In all of these groups, we quantified the percentage of mutations as all alleles different from the reference.

### Differential gene expression analysis from RNA-seq data

Raw reads were processed using the *elvers* (https://github.com/dib-lab/elvers; version 0.1, release DOI: 10.5281/zenodo.3345045) bioinformatic pipeline that utilizes *fastqc* [64], *trimmomatic* [65], and *salmon* [66] to obtain the transcripts per kilobase million (TPM) for each gene. *DESeq2* [67] was used to extract differentially-expressed genes in the Cas9 enzyme or Cas9 mRNA injected samples relative to the uninjected larvae. R package *clusterProfiler* [68] was used to perform enrichment tests of differentially-expressed genes in biological pathways. Network analyses of the common differential expressed genes was performed using the NetworkAnalyst online tool (www.networkanalyst.ca) [69, 70].

### Statistical analyses

All analyses were performed in R version 4.0.2 [71]. Normality of variables was checked using the Shapiro-Wilk test and parametric or nonparametric comparisons made accordingly. Spearman correlation tests (denoted as ρ) and linear regression models were used to determine the relationship between variables. All analyses compared across different experimental batches included *batch* as a factor in the model to prevent biases caused by inter-batch differences. Averages include the standard deviation unless otherwise specified. Alpha to determine significance across the different tests was set at 0.05 unless otherwise specified. Additional R packages used for making figures included *eulerr* [72] and pheatmap [73].

## Abbreviations

CFD: cutting frequency determination
CRISPR: clustered regularly interspaced short palindromic repeats
gRNA: guide RNA
indels: insertions or deletions
PAGE: polyacrylamide gel electrophoresis
RPMN: reads per million normalized

## DECLARATIONS

### Ethics approval and consent to participate

Animal use was approved by the University of California, Davis Institutional Animal Care and Use Committee (IACUC) from the Office of Animal Welfare Assurance accredited by the Association for Assessment and Accreditation of Laboratory Animal Care (Assurance Number A3433-01 on file with the Office of Laboratory Animal Welfare). The IACUC is constituted in accordance with U.S. Public Health Service Animal Welfare Policy and includes a member of the public and a non-scientist. The study and all methods were carried out in accordance with relevant guidelines and regulations and in compliance with the ARRIVE guidelines [74].

### Consent for publication

Not applicable.

### Availability of data and materials

Original fastq files from the CIRCLE-seq and RNA-seq assays are deposited in the European Nucleotide Archive repository under project PRJEB39643.

### Authors’ contributions

JMUS: conceptualization, data collection, analysis, writing of the article. AS, GK, KW, and CI: data collection. MYD: conceptualization, funding acquisition, analysis, writing of the article. All authors have read and approved the manuscript.

## Acknowledgements

We thank our team of UC Davis undergraduate students that maintain husbandry to keep our zebrafish healthy and happy. Thank you to Dr. Li-En Jao for kindly providing the Cas9 mRNA plasmid and comments on the manuscript, and Daniela C. Soto for bioinformatic support. We are grateful to Dr. Charles Vejnar and Dr. Antonio Giraldez for the incorporation of our NHGRIzed zebrafish reference to the CRISPRScan online tool. Finally, we thank the anonymous reviewers of this manuscript for their helpful comments and suggestions that have significantly improved our study.

