## Supplementary Figures for "Evaluation of CRISPR gene-editing tools in zebrafish"

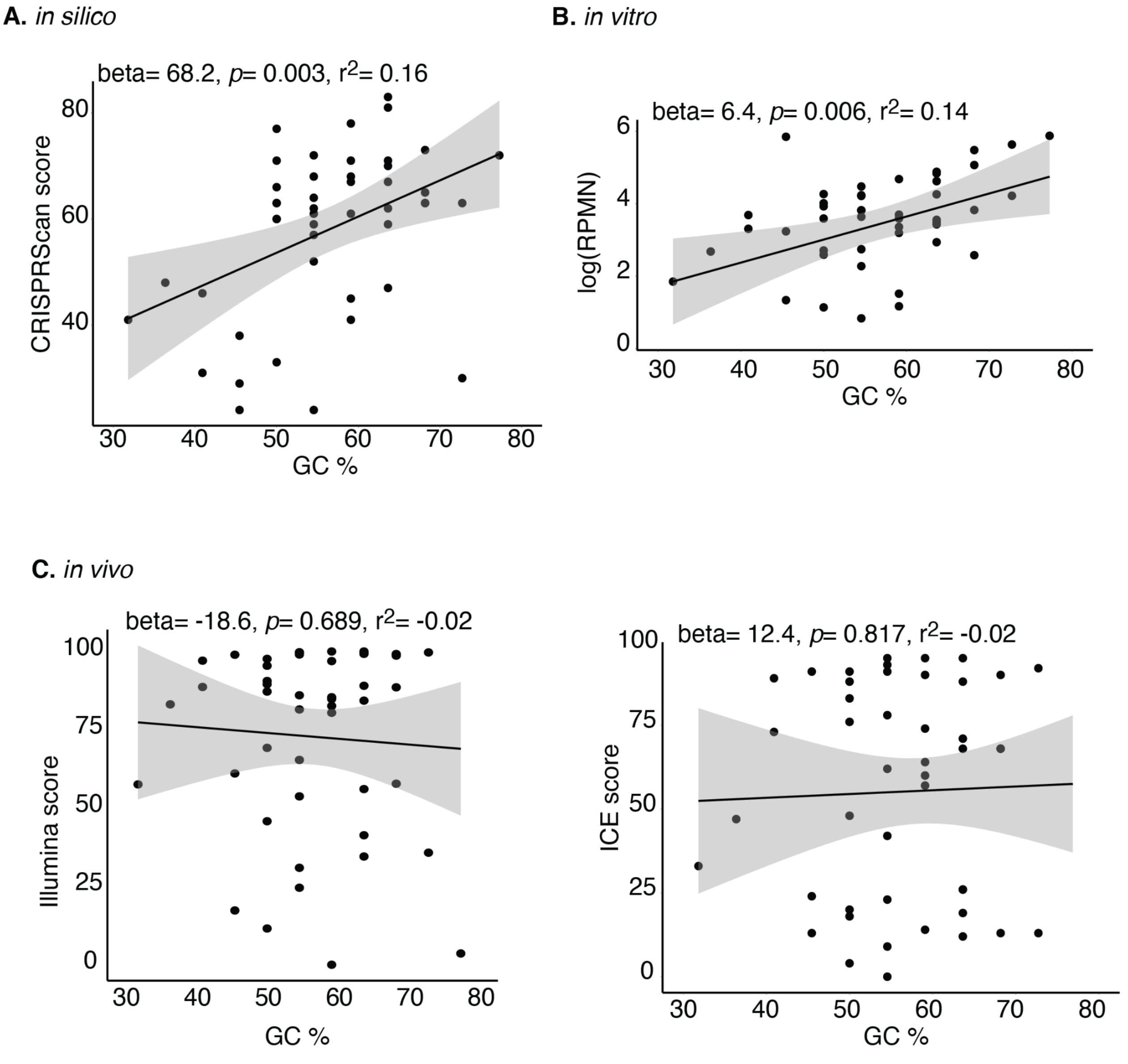


**Supplementary Figure 1. Editing efficiencies across 50 gRNAs with different GC content.** Editing efficiencies **(A)** predicted *in silico* using the CRISPRScan algorithm, **(B)** predicted *in vitro* using the reads per million normalized (RPMN) from the CIRCLE-seq protocol, and **(C)** estimated *in vivo* with Sanger sequencing and the ICE tool (see Methods for details). Results from linear regressions are shown for each comparison.

**Supplementary Figure 2**. **Mosaicism predicted using Illumina sequencing of $G_0$ mutant larvae.** Alleles identified with *CrispRVariants* in larvae injected with Cas9 enzyme ("Cas9enzyme"), Cas9 mRNA ("Cas9RNA"), dCas9 ("dCas9"), scrambled gRNA ("Scrambled"), uninjected batch siblings ("Uninjected"), and a flin-clip from their crossing parents ("Parentals") across 21 genes. Percentages of reads with identified alleles are shown to the right of each allele sequence, with increasing red shades indicating higher percentages.

*actb1*:

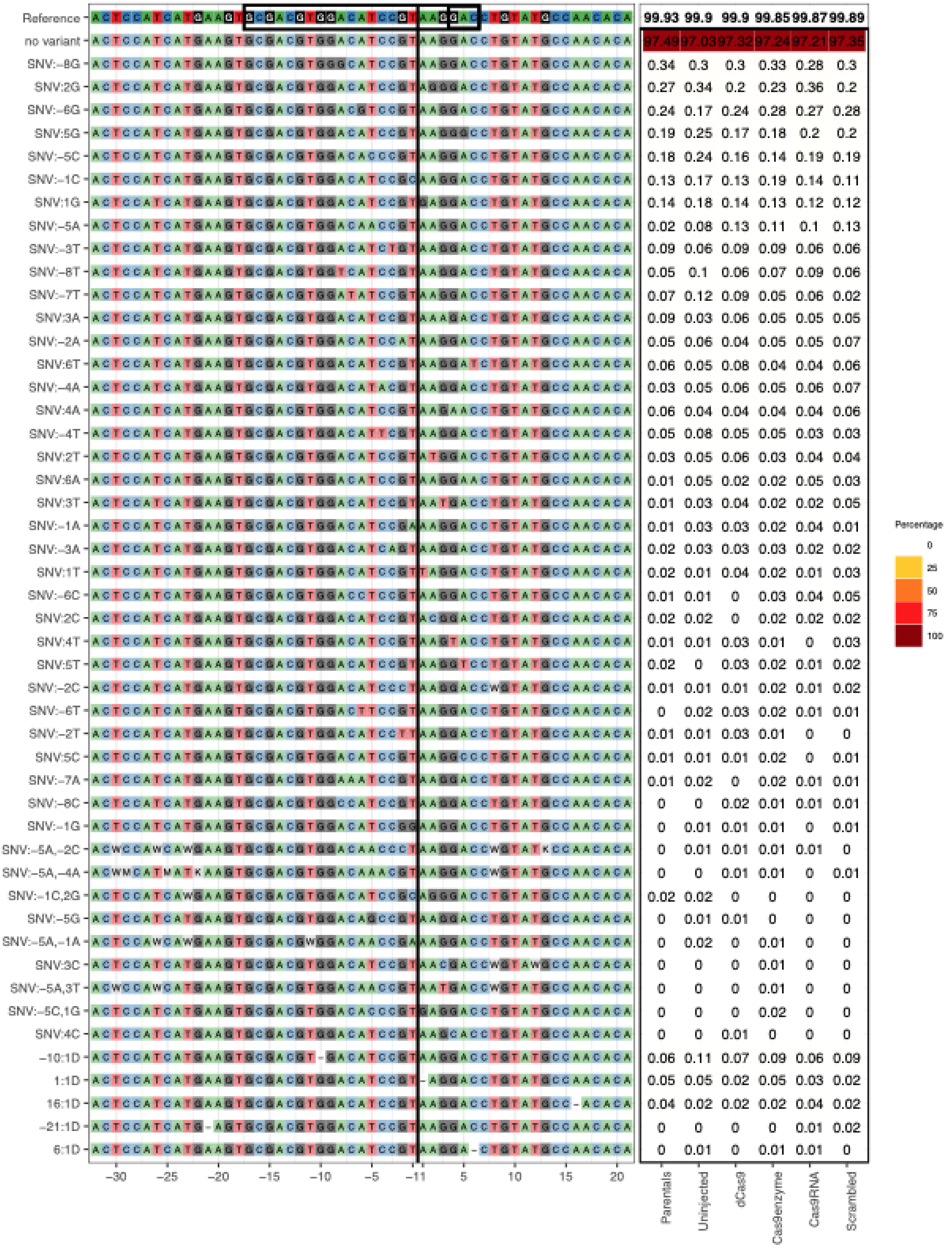

*anp32b*:

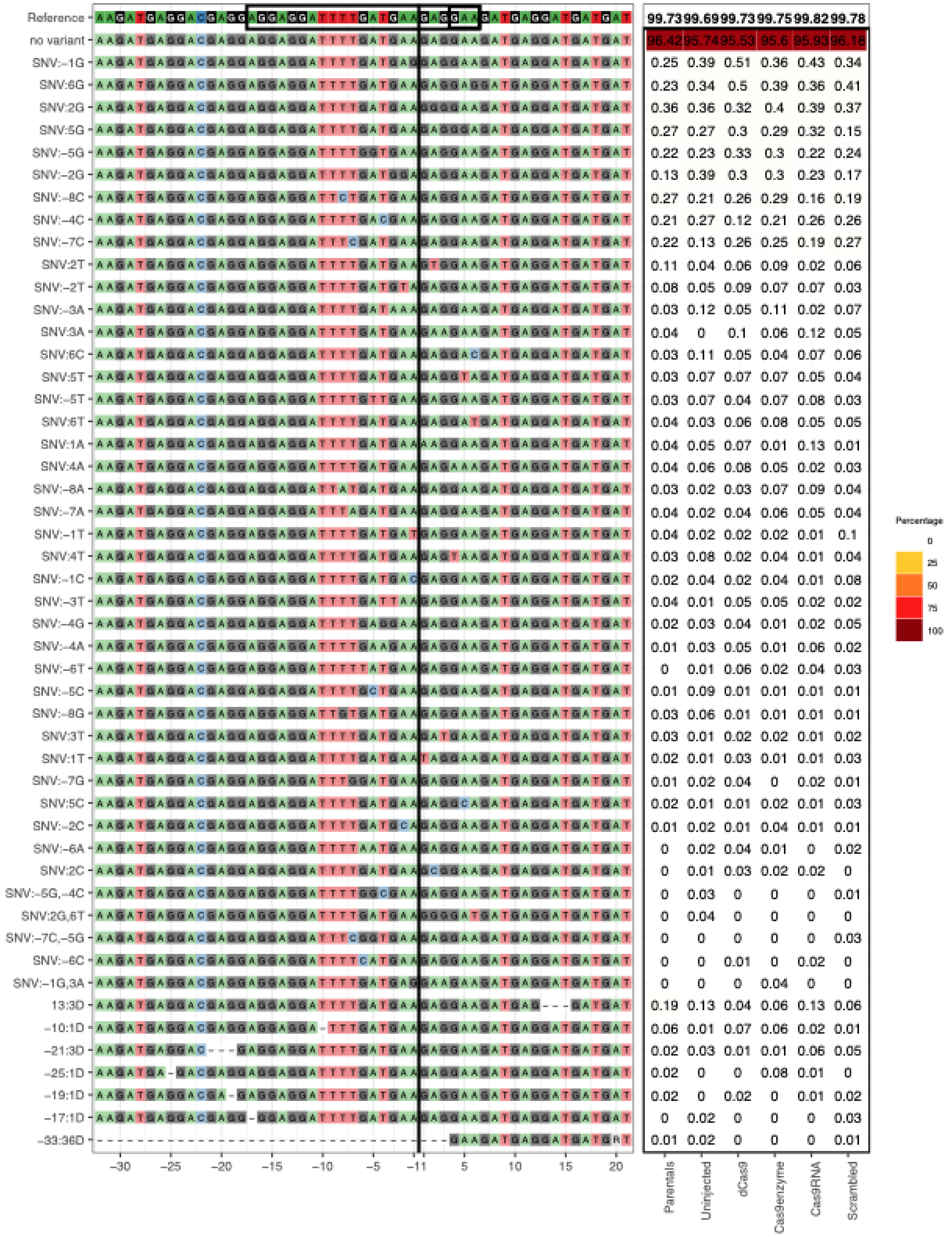

Reference 99.73 99.69 99.73 99.75 99.82 99.78
no variant 96.42 95.74 95.53 95.6 95.93 96.18
SNV:−1G 0.25 0.39 0.51 0.36 0.43 0.34
SNV:6G 0.23 0.34 0.5 0.39 0.36 0.41
SNV:2G 0.36 0.36 0.32 0.4 0.39 0.37
SNV:5G 0.27 0.27 0.3 0.29 0.32 0.15
SNV:−5G 0.22 0.23 0.33 0.3 0.22 0.24
SNV:−2G 0.13 0.39 0.3 0.3 0.23 0.17
SNV:−8C 0.27 0.21 0.26 0.29 0.16 0.19
SNV:−4C 0.21 0.27 0.12 0.21 0.26 0.26
SNV:−7C 0.22 0.13 0.26 0.25 0.19 0.27
SNV:2T 0.11 0.04 0.06 0.09 0.02 0.06
SNV:−2T 0.08 0.05 0.09 0.07 0.07 0.03
SNV:−3A 0.03 0.12 0.05 0.11 0.02 0.07
SNV:3A 0.04 0 0.1 0.06 0.12 0.05
SNV:6C 0.03 0.11 0.05 0.04 0.07 0.06
SNV:5T 0.03 0.07 0.07 0.07 0.05 0.04
SNV:−5T 0.03 0.07 0.04 0.07 0.08 0.03
SNV:6T 0.04 0.03 0.06 0.08 0.05 0.05
SNV:1A 0.04 0.05 0.07 0.01 0.13 0.01
SNV:4A 0.04 0.06 0.08 0.05 0.02 0.03
SNV:−8A 0.03 0.02 0.03 0.07 0.09 0.04
SNV:−7A 0.04 0.02 0.04 0.06 0.05 0.04
SNV:−1T 0.04 0.02 0.02 0.02 0.01 0.1
SNV:4T 0.03 0.08 0.02 0.04 0.01 0.04
SNV:−1C 0.02 0.04 0.02 0.04 0.01 0.08
SNV:−3T 0.04 0.01 0.05 0.05 0.02 0.02
SNV:−4G 0.02 0.03 0.04 0.01 0.02 0.05
SNV:−4A 0.01 0.03 0.05 0.01 0.06 0.02
SNV:−6T 0 0.01 0.06 0.02 0.04 0.03
SNV:−5C 0.01 0.09 0.01 0.01 0.01 0.01
SNV:−8G 0.03 0.06 0.01 0.01 0.01 0.01
SNV:3T 0.03 0.01 0.02 0.02 0.01 0.02
SNV:1T 0.02 0.01 0.03 0.01 0.01 0.03
SNV:−7G 0.01 0.02 0.04 0 0.02 0.01
SNV:5C 0.02 0.01 0.01 0.02 0.01 0.03
SNV:−2C 0.01 0.02 0.01 0.04 0.01 0.01
SNV:−6A 0 0.02 0.04 0.01 0 0.02
SNV:2C 0 0.01 0.03 0.02 0.02 0
SNV:−5G,−4C 0 0.03 0 0 0 0.01
SNV:2G,6T 0 0.04 0 0 0 0
SNV:−7C,−5G 0 0 0 0 0 0.03
SNV:−6C 0 0 0.01 0 0.02 0
SNV:−1G,3A 0 0 0 0.04 0 0
13:3D 0.19 0.13 0.04 0.06 0.13 0.06
−10:1D 0.06 0.01 0.07 0.06 0.02 0.01
−21:3D 0.02 0.03 0.01 0.01 0.06 0.05
−25:1D 0.02 0 0 0.08 0.01 0
−19:1D 0.02 0 0.02 0 0.01 0.02
−17:1D 0 0.02 0 0 0 0.03
−33:36D 0.01 0.02 0 0 0 0.01
−30 −25 −20 −15 −10 −5 −1 1 5 10 15 20
Parentals
Uninjected
dCas9
Cas9enzyme
Cas9RNA
Scrambled
Percentage
0
25
50
75
100

*atp1a1a.1*:

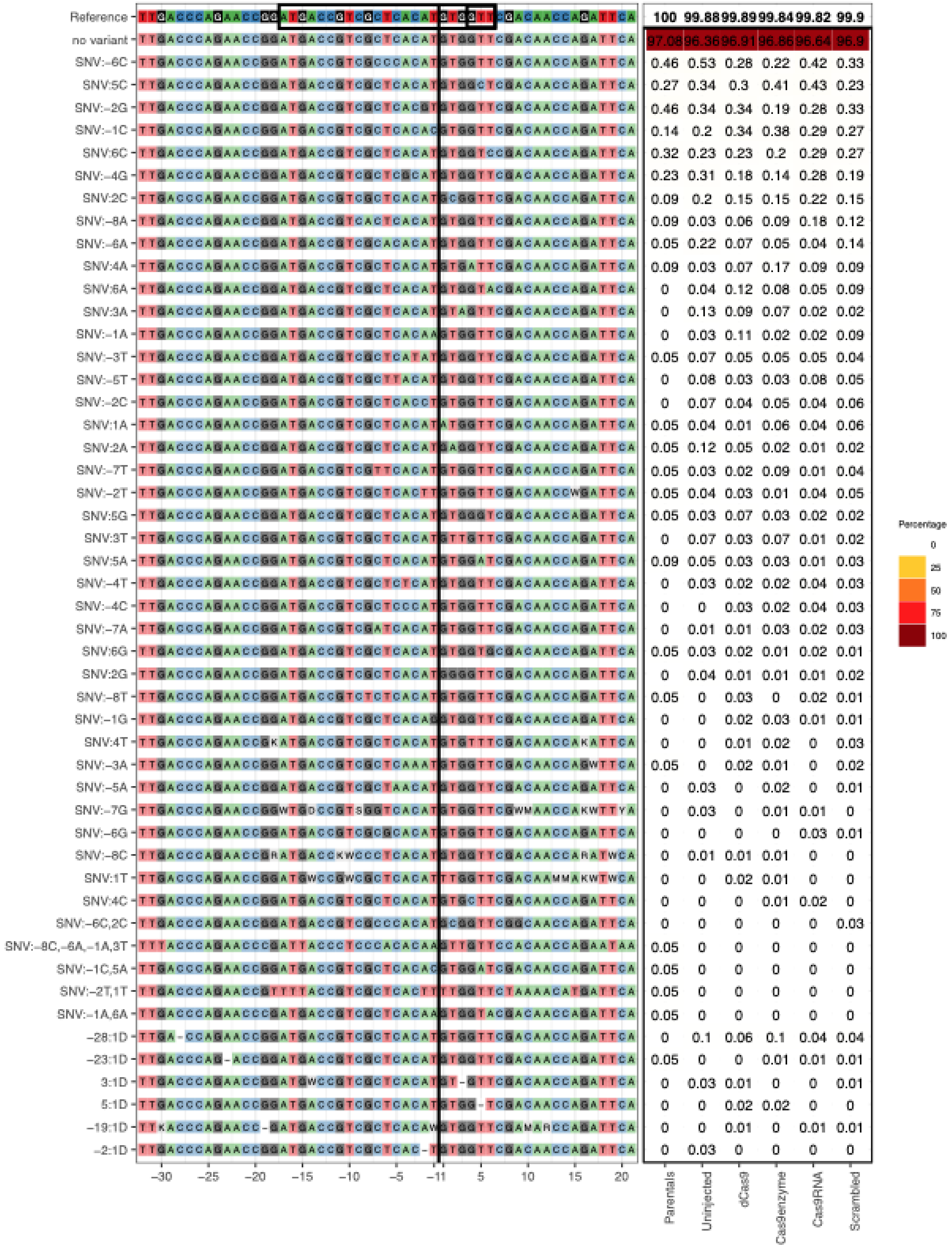

Reference 100 99.88 99.89 99.84 99.82 99.9
no variant 97.08 96.36 96.91 96.86 96.64 96.9
SNV:-6C 0.46 0.53 0.28 0.22 0.42 0.33
SNV:5C 0.27 0.34 0.3 0.41 0.43 0.23
SNV:-2G 0.46 0.34 0.34 0.19 0.28 0.33
SNV:-1C 0.14 0.2 0.34 0.38 0.29 0.27
SNV:6C 0.32 0.23 0.23 0.2 0.29 0.27
SNV:-4G 0.23 0.31 0.18 0.14 0.28 0.19
SNV:2C 0.09 0.2 0.15 0.15 0.22 0.15
SNV:-8A 0.09 0.03 0.06 0.09 0.18 0.12
SNV:-6A 0.05 0.22 0.07 0.05 0.04 0.14
SNV:4A 0.09 0.03 0.07 0.17 0.09 0.09
SNV:6A 0 0.04 0.12 0.08 0.05 0.09
SNV:3A 0 0.13 0.09 0.07 0.02 0.02
SNV:-1A 0 0.03 0.11 0.02 0.02 0.09
SNV:-3T 0.05 0.07 0.05 0.05 0.05 0.04
SNV:-5T 0 0.08 0.03 0.03 0.08 0.05
SNV:-2C 0 0.07 0.04 0.05 0.04 0.06
SNV:1A 0.05 0.04 0.01 0.06 0.04 0.06
SNV:2A 0.05 0.12 0.05 0.02 0.01 0.02
SNV:-7T 0.05 0.03 0.02 0.09 0.01 0.04
SNV:-2T 0.05 0.04 0.03 0.01 0.04 0.05
SNV:5G 0.05 0.03 0.07 0.03 0.02 0.02
SNV:3T 0 0.07 0.03 0.07 0.01 0.02
SNV:5A 0.09 0.05 0.03 0.03 0.01 0.03
SNV:-4T 0 0.03 0.02 0.02 0.04 0.03
SNV:-4C 0 0 0.03 0.02 0.04 0.03
SNV:-7A 0 0.01 0.01 0.03 0.02 0.03
SNV:6G 0.05 0.03 0.02 0.01 0.02 0.01
SNV:2G 0 0.04 0.01 0.01 0.01 0.02
SNV:-8T 0.05 0 0.03 0 0.02 0.01
SNV:-1G 0 0 0.02 0.03 0.01 0.01
SNV:4T 0 0 0.01 0.02 0 0.03
SNV:-3A 0.05 0 0.02 0.01 0 0.02
SNV:-5A 0 0.03 0 0.02 0 0.01
SNV:-7G 0 0.03 0 0.01 0.01 0
SNV:-6G 0 0 0 0 0.03 0.01
SNV:-8C 0 0.01 0.01 0.01 0 0
SNV:1T 0 0 0.02 0.01 0 0
SNV:4C 0 0 0 0.01 0.02 0
SNV:-6C,2C 0 0 0 0 0 0.03
SNV:-8C,-6A,-1A,3T 0.05 0 0 0 0 0
SNV:-1C,5A 0.05 0 0 0 0 0
SNV:-2T,1T 0.05 0 0 0 0 0
SNV:-1A,6A 0.05 0 0 0 0 0
-28:1D 0 0.1 0.06 0.1 0.04 0.04
-23:1D 0.05 0 0 0.01 0.01 0.01
3:1D 0 0.03 0.01 0 0 0.01
5:1D 0 0 0.02 0.02 0 0
-19:1D 0 0 0.01 0 0.01 0.01
-2:1D 0 0.03 0 0 0 0
-30 -25 -20 -15 -10 -5 -1 1 5 10 15 20
Parentals
Uninjected
dCas9
Cas9enzyme
Cas9RNA
Scrambled
Percentage
0
25
50
75
100

*atp5md*:

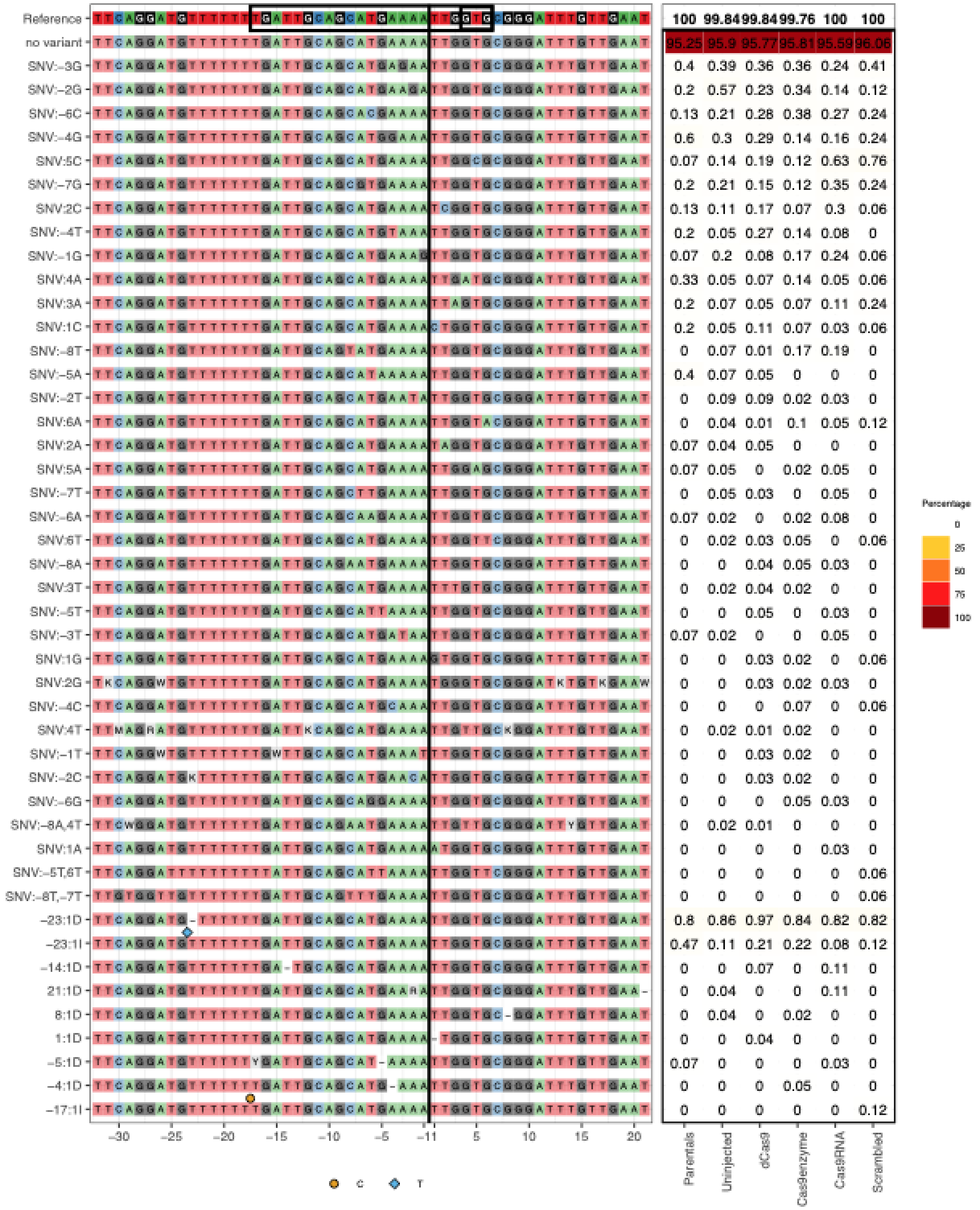
Reference 100 99.84 99.84 99.76 100 100
no variant 95.25 95.9 95.77 95.81 95.59 96.06
SNV:-3G 0.4 0.39 0.36 0.36 0.24 0.41
SNV:-2G 0.2 0.57 0.23 0.34 0.14 0.12
SNV:-6C 0.13 0.21 0.28 0.38 0.27 0.24
SNV:-4G 0.6 0.3 0.29 0.14 0.16 0.24
SNV:5C 0.07 0.14 0.19 0.12 0.63 0.76
SNV:-7G 0.2 0.21 0.15 0.12 0.35 0.24
SNV:2C 0.13 0.11 0.17 0.07 0.3 0.06
SNV:-4T 0.2 0.05 0.27 0.14 0.08 0
SNV:-1G 0.07 0.2 0.08 0.17 0.24 0.06
SNV:4A 0.33 0.05 0.07 0.14 0.05 0.06
SNV:3A 0.2 0.07 0.05 0.07 0.11 0.24
SNV:1C 0.2 0.05 0.11 0.07 0.03 0.06
SNV:-8T 0 0.07 0.01 0.17 0.19 0
SNV:-5A 0.4 0.07 0.05 0 0 0
SNV:-2T 0 0.09 0.09 0.02 0.03 0
SNV:6A 0 0.04 0.01 0.1 0.05 0.12
SNV:2A 0.07 0.04 0.05 0 0 0
SNV:5A 0.07 0.05 0 0.02 0.05 0
SNV:-7T 0 0.05 0.03 0 0.05 0
SNV:-6A 0.07 0.02 0 0.02 0.08 0
SNV:6T 0 0.02 0.03 0.05 0 0.06
SNV:-8A 0 0 0.04 0.05 0.03 0
SNV:3T 0 0.02 0.04 0.02 0 0
SNV:-5T 0 0 0.05 0 0.03 0
SNV:-3T 0.07 0.02 0 0 0.05 0
SNV:1G 0 0 0.03 0.02 0 0.06
SNV:2G 0 0 0.03 0.02 0.03 0
SNV:-4C 0 0 0 0.07 0 0.06
SNV:4T 0 0.02 0.01 0.02 0 0
SNV:-1T 0 0 0.03 0.02 0 0
SNV:-2C 0 0 0.03 0.02 0 0
SNV:-6G 0 0 0 0.05 0.03 0
SNV:-8A,4T 0 0.02 0.01 0 0 0
SNV:1A 0 0 0 0 0.03 0
SNV:-5T,6T 0 0 0 0 0 0.06
SNV:-8T,-7T 0 0 0 0 0 0.06
-23:1D 0.8 0.86 0.97 0.84 0.82 0.82
-23:1I 0.47 0.11 0.21 0.22 0.08 0.12
-14:1D 0 0 0.07 0 0.11 0
21:1D 0 0.04 0 0 0.11 0
8:1D 0 0.04 0 0.02 0 0
1:1D 0 0 0.04 0 0 0
-5:1D 0.07 0 0 0 0.03 0
-4:1D 0 0 0 0.05 0 0
-17:1I 0 0 0 0 0 0.12
-30 -25 -20 -15 -10 -5 -1 1 5 10 15 20
Parentals
Uninjected
dCas9
Cas9enzyme
Cas9RNA
Scrambled
Percentage
0
25
50
75
100
C
T

*c3a.1:*

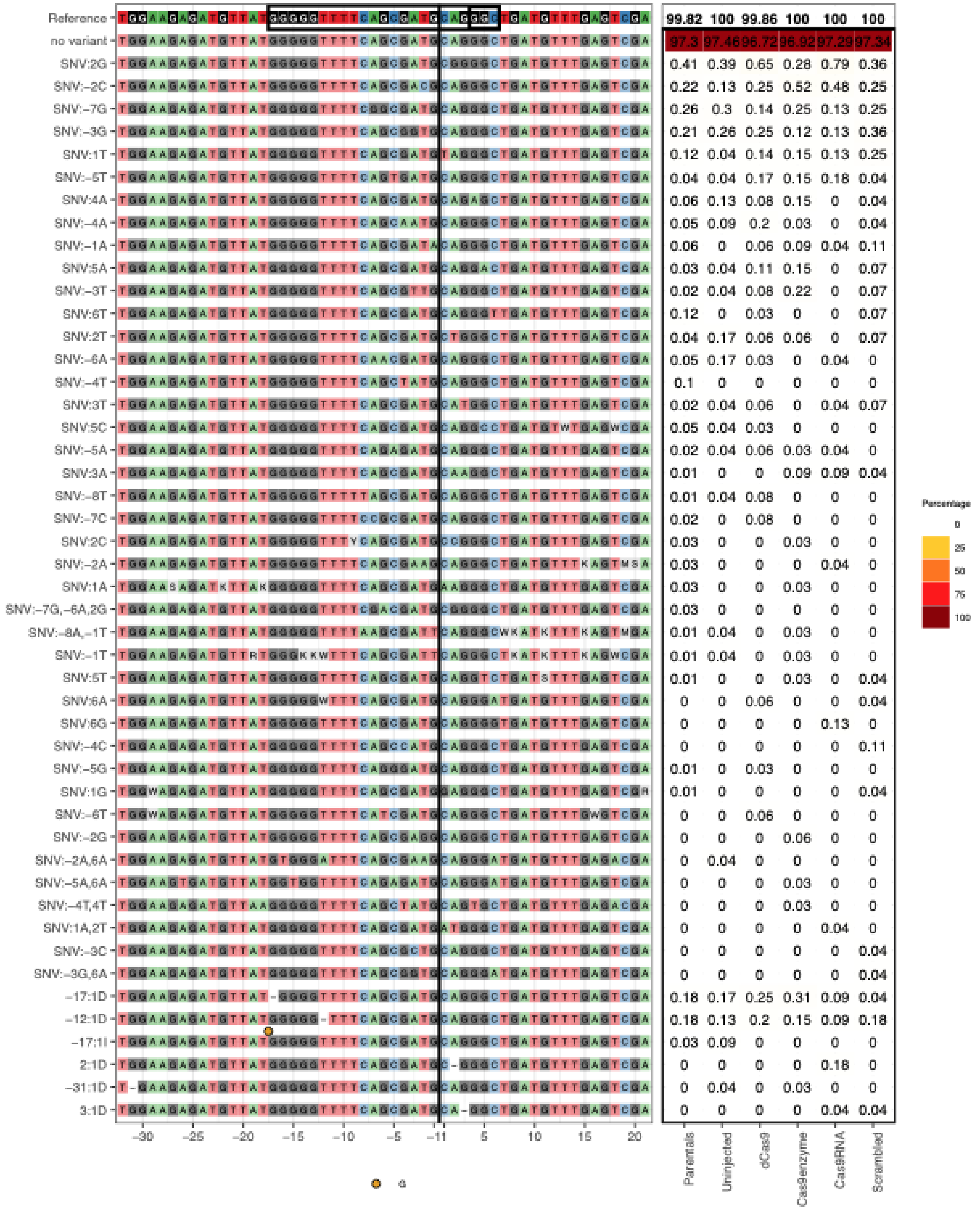

Reference 99.82 100 99.86 100 100 100
no variant 97.3 97.46 96.72 96.92 97.29 97.34
SNV:2G 0.41 0.39 0.65 0.28 0.79 0.36
SNV:−2C 0.22 0.13 0.25 0.52 0.48 0.25
SNV:−7G 0.26 0.3 0.14 0.25 0.13 0.25
SNV:−3G 0.21 0.26 0.25 0.12 0.13 0.36
SNV:1T 0.12 0.04 0.14 0.15 0.13 0.25
SNV:−5T 0.04 0.04 0.17 0.15 0.18 0.04
SNV:4A 0.06 0.13 0.08 0.15 0 0.04
SNV:−4A 0.05 0.09 0.2 0.03 0 0.04
SNV:−1A 0.06 0 0.06 0.09 0.04 0.11
SNV:5A 0.03 0.04 0.11 0.15 0 0.07
SNV:−3T 0.02 0.04 0.08 0.22 0 0.07
SNV:6T 0.12 0 0.03 0 0 0.07
SNV:2T 0.04 0.17 0.06 0.06 0 0.07
SNV:−6A 0.05 0.17 0.03 0 0.04 0
SNV:−4T 0.1 0 0 0 0 0
SNV:3T 0.02 0.04 0.06 0 0.04 0.07
SNV:5C 0.05 0.04 0.03 0 0 0
SNV:−5A 0.02 0.04 0.06 0.03 0.04 0
SNV:3A 0.01 0 0 0.09 0.09 0.04
SNV:−8T 0.01 0.04 0.08 0 0 0
SNV:−7C 0.02 0 0.08 0 0 0
SNV:2C 0.03 0 0 0.03 0 0
SNV:−2A 0.03 0 0 0 0.04 0
SNV:1A 0.03 0 0 0.03 0 0
SNV:−7G,−6A,2G 0.03 0 0 0 0 0
SNV:−8A,−1T 0.01 0.04 0 0.03 0 0
SNV:−1T 0.01 0.04 0 0.03 0 0
SNV:5T 0.01 0 0 0.03 0 0.04
SNV:6A 0 0 0.06 0 0 0.04
SNV:6G 0 0 0 0 0.13 0
SNV:−4C 0 0 0 0 0 0.11
SNV:−5G 0.01 0 0.03 0 0 0
SNV:1G 0.01 0 0 0 0 0.04
SNV:−6T 0 0 0.06 0 0 0
SNV:−2G 0 0 0 0.06 0 0
SNV:−2A,6A 0 0.04 0 0 0 0
SNV:−5A,6A 0 0 0 0.03 0 0
SNV:−4T,4T 0 0 0 0.03 0 0
SNV:1A,2T 0 0 0 0 0.04 0
SNV:−3C 0 0 0 0 0 0.04
SNV:−3G,6A 0 0 0 0 0 0.04
−17:1D 0.18 0.17 0.25 0.31 0.09 0.04
−12:1D 0.18 0.13 0.2 0.15 0.09 0.18
−17:1I 0.03 0.09 0 0 0 0
2:1D 0 0 0 0 0.18 0
−31:1D 0 0.04 0 0.03 0 0
3:1D 0 0 0 0 0.04 0.04
−30 −25 −20 −15 −10 −5 −1 1 5 10 15 20
Parentals
Uninjected
dCas9
Cas9enzyme
Cas9RNA
Scrambled
G
Percentage
0
25
50
75
100

*fabp2*:

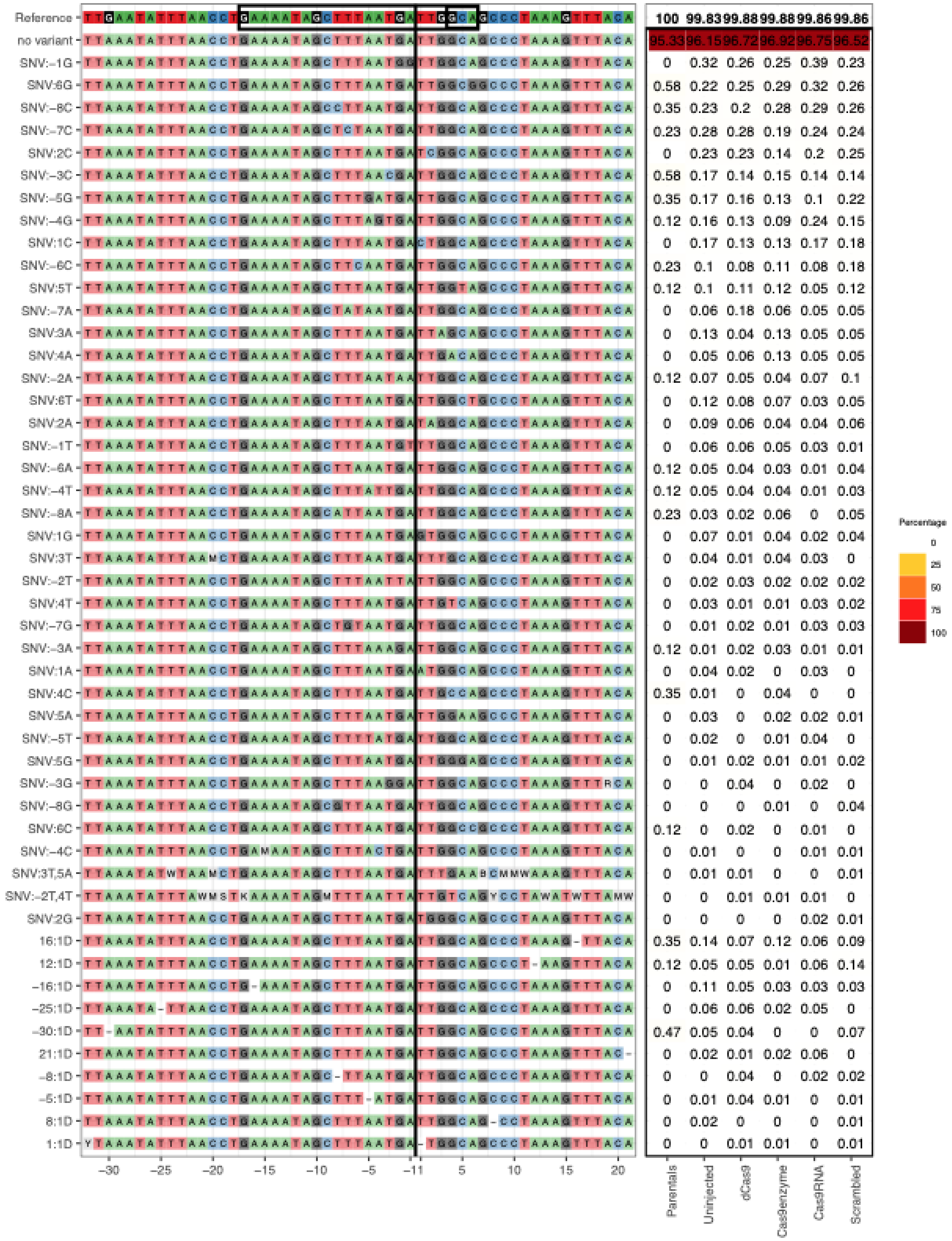
Reference 100 99.83 99.88 99.88 99.86 99.86
no variant 95.33 96.15 96.72 96.92 96.75 96.52
SNV:-1G 0 0.32 0.26 0.25 0.39 0.23
SNV:6G 0.58 0.22 0.25 0.29 0.32 0.26
SNV:-8C 0.35 0.23 0.2 0.28 0.29 0.26
SNV:-7C 0.23 0.28 0.28 0.19 0.24 0.24
SNV:2C 0 0.23 0.23 0.14 0.2 0.25
SNV:-3C 0.58 0.17 0.14 0.15 0.14 0.14
SNV:-5G 0.35 0.17 0.16 0.13 0.1 0.22
SNV:-4G 0.12 0.16 0.13 0.09 0.24 0.15
SNV:1C 0 0.17 0.13 0.13 0.17 0.18
SNV:-6C 0.23 0.1 0.08 0.11 0.08 0.18
SNV:5T 0.12 0.1 0.11 0.12 0.05 0.12
SNV:-7A 0 0.06 0.18 0.06 0.05 0.05
SNV:3A 0 0.13 0.04 0.13 0.05 0.05
SNV:4A 0 0.05 0.06 0.13 0.05 0.05
SNV:-2A 0.12 0.07 0.05 0.04 0.07 0.1
SNV:6T 0 0.12 0.08 0.07 0.03 0.05
SNV:2A 0 0.09 0.06 0.04 0.04 0.06
SNV:-1T 0 0.06 0.06 0.05 0.03 0.01
SNV:-6A 0.12 0.05 0.04 0.03 0.01 0.04
SNV:-4T 0.12 0.05 0.04 0.04 0.01 0.03
SNV:-8A 0.23 0.03 0.02 0.06 0 0.05
SNV:1G 0 0.07 0.01 0.04 0.02 0.04
SNV:3T 0 0.04 0.01 0.04 0.03 0
SNV:-2T 0 0.02 0.03 0.02 0.02 0.02
SNV:4T 0 0.03 0.01 0.01 0.03 0.02
SNV:-7G 0 0.01 0.02 0.01 0.03 0.03
SNV:-3A 0.12 0.01 0.02 0.03 0.01 0.01
SNV:1A 0 0.04 0.02 0 0.03 0
SNV:4C 0.35 0.01 0 0.04 0 0
SNV:5A 0 0.03 0 0.02 0.02 0.01
SNV:-5T 0 0.02 0 0.01 0.04 0
SNV:5G 0 0.01 0.02 0.01 0.01 0.02
SNV:-3G 0 0 0.04 0 0.02 0
SNV:-8G 0 0 0 0.01 0 0.04
SNV:6C 0.12 0 0.02 0 0.01 0
SNV:-4C 0 0.01 0 0 0.01 0.01
SNV:3T,5A 0 0.01 0.01 0 0 0.01
SNV:-2T,4T 0 0 0.01 0.01 0.01 0
SNV:2G 0 0 0 0 0.02 0.01
16:1D 0.35 0.14 0.07 0.12 0.06 0.09
12:1D 0.12 0.05 0.05 0.01 0.06 0.14
-16:1D 0 0.11 0.05 0.03 0.03 0.03
-25:1D 0 0.06 0.06 0.02 0.05 0
-30:1D 0.47 0.05 0.04 0 0 0.07
21:1D 0 0.02 0.01 0.02 0.06 0
-8:1D 0 0 0.04 0 0.02 0.02
-5:1D 0 0.01 0.04 0.01 0 0.01
8:1D 0 0.02 0 0 0 0.01
1:1D 0 0 0.01 0.01 0 0.01
-30 -25 -20 -15 -10 -5 -1 1 5 10 15 20
Parentals
Uninjected
dCas9
Cas9enzyme
Cas9RNA
Scrambled
Percentage
0
25
50
75
100

*fga*:

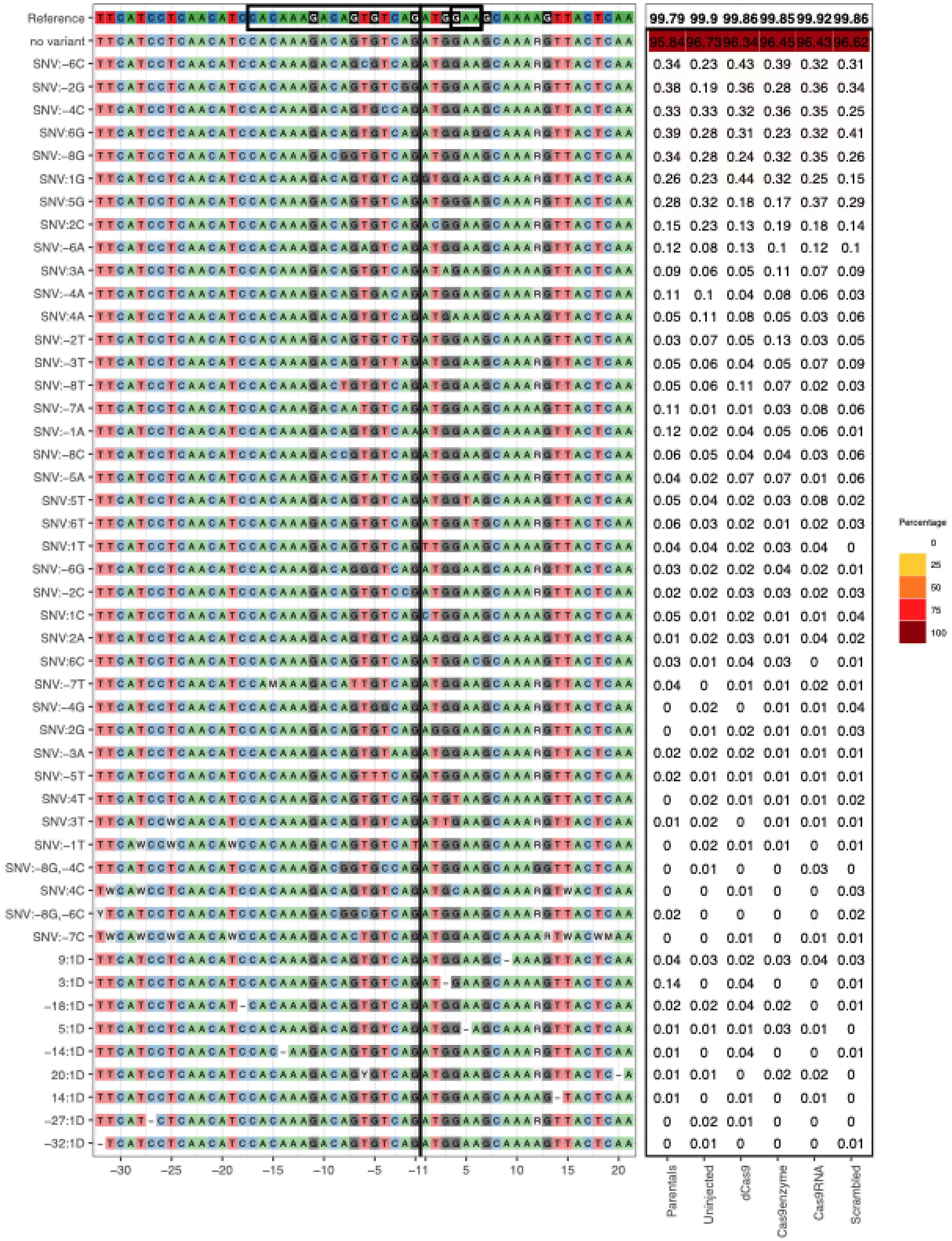
Reference 99.79 99.9 99.86 99.85 99.92 99.86
no variant 95.84 96.73 96.34 96.45 96.43 96.62
SNV:-6C 0.34 0.23 0.43 0.39 0.32 0.31
SNV:-2G 0.38 0.19 0.36 0.28 0.36 0.34
SNV:-4C 0.33 0.33 0.32 0.36 0.35 0.25
SNV:6G 0.39 0.28 0.31 0.23 0.32 0.41
SNV:-8G 0.34 0.28 0.24 0.32 0.35 0.26
SNV:1G 0.26 0.23 0.44 0.32 0.25 0.15
SNV:5G 0.28 0.32 0.18 0.17 0.37 0.29
SNV:2C 0.15 0.23 0.13 0.19 0.18 0.14
SNV:-6A 0.12 0.08 0.13 0.1 0.12 0.1
SNV:3A 0.09 0.06 0.05 0.11 0.07 0.09
SNV:-4A 0.11 0.1 0.04 0.08 0.06 0.03
SNV:4A 0.05 0.11 0.08 0.05 0.03 0.06
SNV:-2T 0.03 0.07 0.05 0.13 0.03 0.05
SNV:-3T 0.05 0.06 0.04 0.05 0.07 0.09
SNV:-8T 0.05 0.06 0.11 0.07 0.02 0.03
SNV:-7A 0.11 0.01 0.01 0.03 0.08 0.06
SNV:-1A 0.12 0.02 0.04 0.05 0.06 0.01
SNV:-8C 0.06 0.05 0.04 0.04 0.03 0.06
SNV:-5A 0.04 0.02 0.07 0.07 0.01 0.06
SNV:5T 0.05 0.04 0.02 0.03 0.08 0.02
SNV:6T 0.06 0.03 0.02 0.01 0.02 0.03
SNV:1T 0.04 0.04 0.02 0.03 0.04 0
SNV:-6G 0.03 0.02 0.02 0.04 0.02 0.01
SNV:-2C 0.02 0.02 0.03 0.03 0.02 0.03
SNV:1C 0.05 0.01 0.02 0.01 0.01 0.04
SNV:2A 0.01 0.02 0.03 0.01 0.04 0.02
SNV:6C 0.03 0.01 0.04 0.03 0 0.01
SNV:-7T 0.04 0 0.01 0.01 0.02 0.01
SNV:-4G 0 0.02 0 0.01 0.01 0.04
SNV:2G 0 0.01 0.02 0.01 0.01 0.03
SNV:-3A 0.02 0.02 0.02 0.01 0.01 0.01
SNV:-5T 0.02 0.01 0.01 0.01 0.01 0.01
SNV:4T 0 0.02 0.01 0.01 0.01 0.02
SNV:3T 0.01 0.02 0 0.01 0.01 0.01
SNV:-1T 0 0.02 0.01 0.01 0 0.01
SNV:-8G,-4C 0 0.01 0 0 0.03 0
SNV:4C 0 0 0.01 0 0 0.03
SNV:-8G,-6C 0.02 0 0 0 0 0.02
SNV:-7C 0 0 0.01 0 0.01 0.01
9:1D 0.04 0.03 0.02 0.03 0.04 0.03
3:1D 0.14 0 0.04 0 0 0.01
-18:1D 0.02 0.02 0.04 0.02 0 0.01
5:1D 0.01 0.01 0.01 0.03 0.01 0
-14:1D 0.01 0 0.04 0 0 0.01
20:1D 0.01 0.01 0 0.02 0.02 0
14:1D 0.01 0 0.01 0 0.01 0
-27:1D 0 0.02 0.01 0 0 0
-32:1D 0 0.01 0 0 0 0.01
-30 -25 -20 -15 -10 -5 -1 1 5 10 15 20
Parentals
Uninjected
dCas9
Cas9enzyme
Cas9RNA
Scrambled
Percentage
0
25
50
75
100

*fgg*:

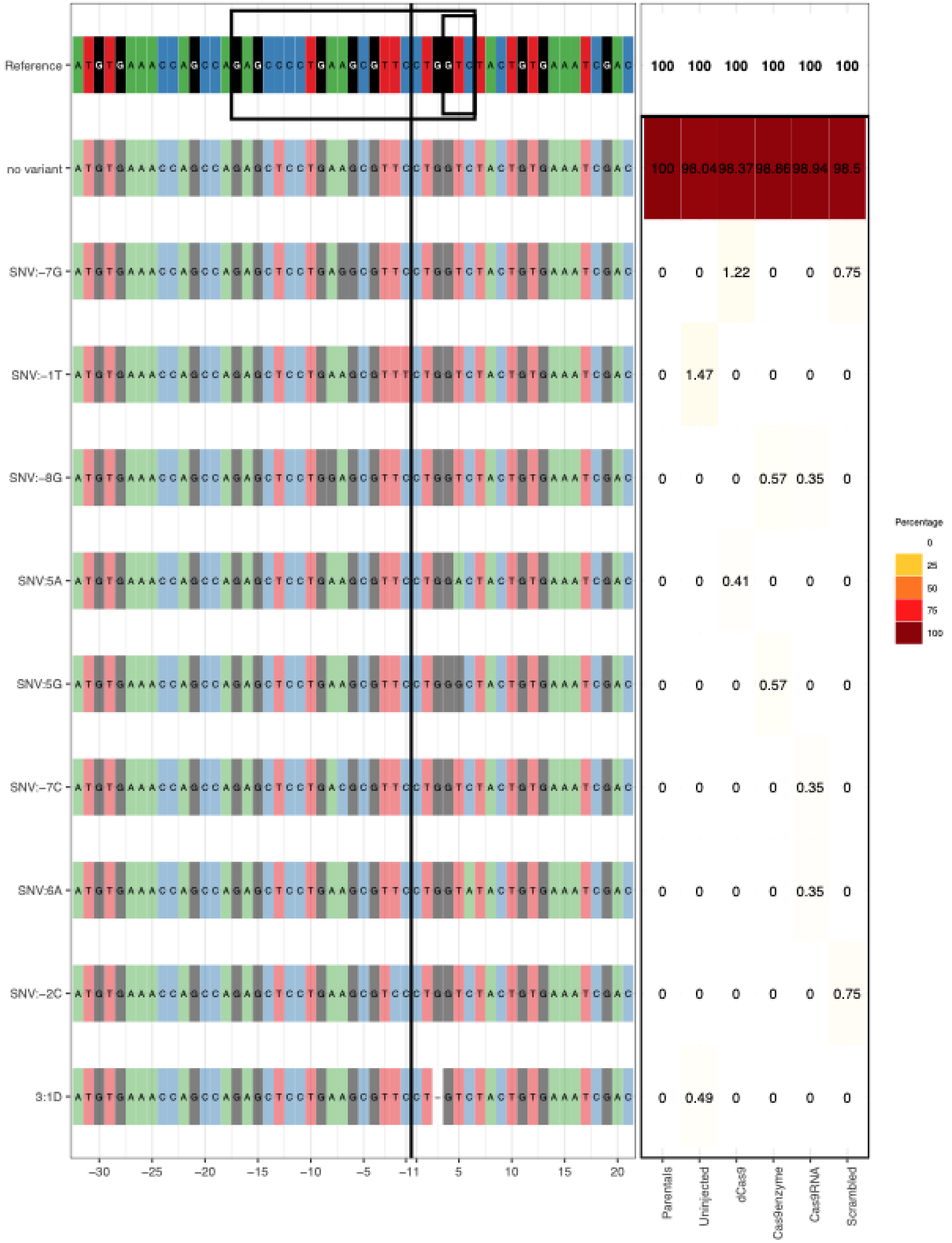

Reference
no variant
SNV:−7G
SNV:−1T
SNV:−8G
SNV:5A
SNV:5G
SNV:−7C
SNV:6A
SNV:−2C
3:1D
100 100 100 100 100 100
100 98.04 98.37 98.86 98.94 98.5
0 0 1.22 0 0 0.75
0 1.47 0 0 0 0
0 0 0 0.57 0.35 0
0 0 0.41 0 0 0
0 0 0 0.57 0 0
0 0 0 0 0.35 0
0 0 0 0 0.35 0
0 0 0 0 0 0.75
0 0.49 0 0 0 0
−30 −25 −20 −15 −10 −5 −1 1 5 10 15 20
Parentals
Uninjected
dCas9
Cas9enzyme
Cas9RNA
Scrambled
Percentage
0
25
50
75
100

*galnt7*:

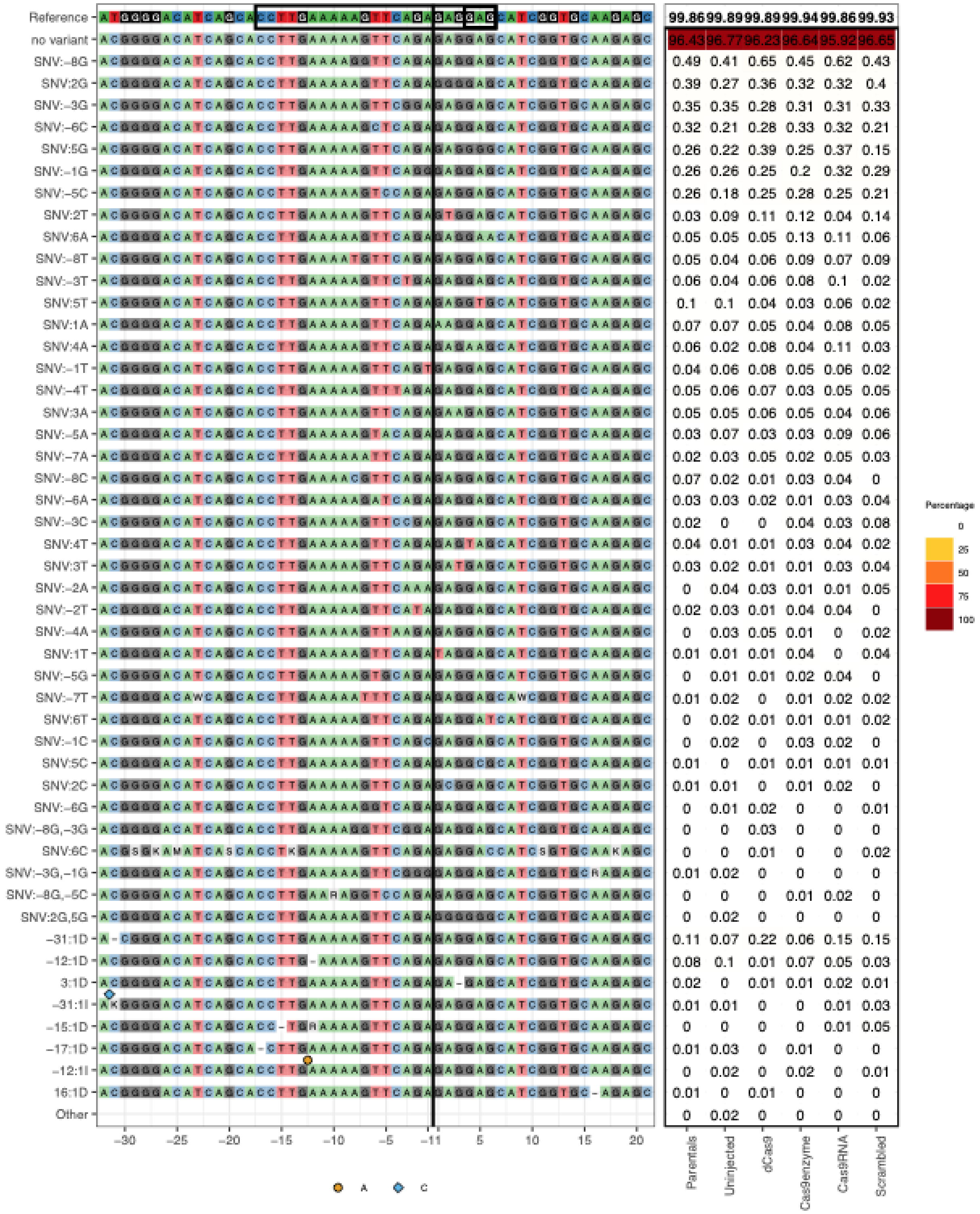
Reference 99.86 99.89 99.89 99.94 99.86 99.93
no variant 96.43 96.77 96.23 96.64 95.92 96.65
SNV:−8G 0.49 0.41 0.65 0.45 0.62 0.43
SNV:2G 0.39 0.27 0.36 0.32 0.32 0.4
SNV:−3G 0.35 0.35 0.28 0.31 0.31 0.33
SNV:−6C 0.32 0.21 0.28 0.33 0.32 0.21
SNV:5G 0.26 0.22 0.39 0.25 0.37 0.15
SNV:−1G 0.26 0.26 0.25 0.2 0.32 0.29
SNV:−5C 0.26 0.18 0.25 0.28 0.25 0.21
SNV:2T 0.03 0.09 0.11 0.12 0.04 0.14
SNV:6A 0.05 0.05 0.05 0.13 0.11 0.06
SNV:−8T 0.05 0.04 0.06 0.09 0.07 0.09
SNV:−3T 0.06 0.04 0.06 0.08 0.1 0.02
SNV:5T 0.1 0.1 0.04 0.03 0.06 0.02
SNV:1A 0.07 0.07 0.05 0.04 0.08 0.05
SNV:4A 0.06 0.02 0.08 0.04 0.11 0.03
SNV:−1T 0.04 0.06 0.08 0.05 0.06 0.02
SNV:−4T 0.05 0.06 0.07 0.03 0.05 0.05
SNV:3A 0.05 0.05 0.06 0.05 0.04 0.06
SNV:−5A 0.03 0.07 0.03 0.03 0.09 0.06
SNV:−7A 0.02 0.03 0.05 0.02 0.05 0.03
SNV:−8C 0.07 0.02 0.01 0.03 0.04 0
SNV:−6A 0.03 0.03 0.02 0.01 0.03 0.04
SNV:−3C 0.02 0 0 0.04 0.03 0.08
SNV:4T 0.04 0.01 0.01 0.03 0.04 0.02
SNV:3T 0.03 0.02 0.01 0.01 0.03 0.04
SNV:−2A 0 0.04 0.03 0.01 0.01 0.05
SNV:−2T 0.02 0.03 0.01 0.04 0.04 0
SNV:−4A 0 0.03 0.05 0.01 0 0.02
SNV:1T 0.01 0.01 0.01 0.04 0 0.04
SNV:−5G 0 0.01 0.01 0.02 0.04 0
SNV:−7T 0.01 0.02 0 0.01 0.02 0.02
SNV:6T 0 0.02 0.01 0.01 0.01 0.02
SNV:−1C 0 0.02 0 0.03 0.02 0
SNV:5C 0.01 0 0.01 0.01 0.01 0.01
SNV:2C 0.01 0.01 0 0.01 0.02 0
SNV:−6G 0 0.01 0.02 0 0 0.01
SNV:−8G,−3G 0 0 0.03 0 0 0
SNV:6C 0 0 0.01 0 0 0.02
SNV:−3G,−1G 0.01 0.02 0 0 0 0
SNV:−8G,−5C 0 0 0 0.01 0.02 0
SNV:2G,5G 0 0.02 0 0 0 0
−31:1D 0.11 0.07 0.22 0.06 0.15 0.15
−12:1D 0.08 0.1 0.01 0.07 0.05 0.03
3:1D 0.02 0 0.01 0.01 0.02 0.01
−31:1I 0.01 0.01 0 0 0.01 0.03
−15:1D 0 0 0 0 0.01 0.05
−17:1D 0.01 0.03 0 0.01 0 0
−12:1I 0 0.02 0 0.02 0 0.01
16:1D 0.01 0 0.01 0 0 0
Other 0 0.02 0 0 0 0
−30 −25 −20 −15 −10 −5 −1 1 5 10 15 20
Parentals Uninjected dCas9 Cas9enzyme Cas9RNA Scrambled
A C
Percentage
0 25 50 75 100

*ggact.2*:

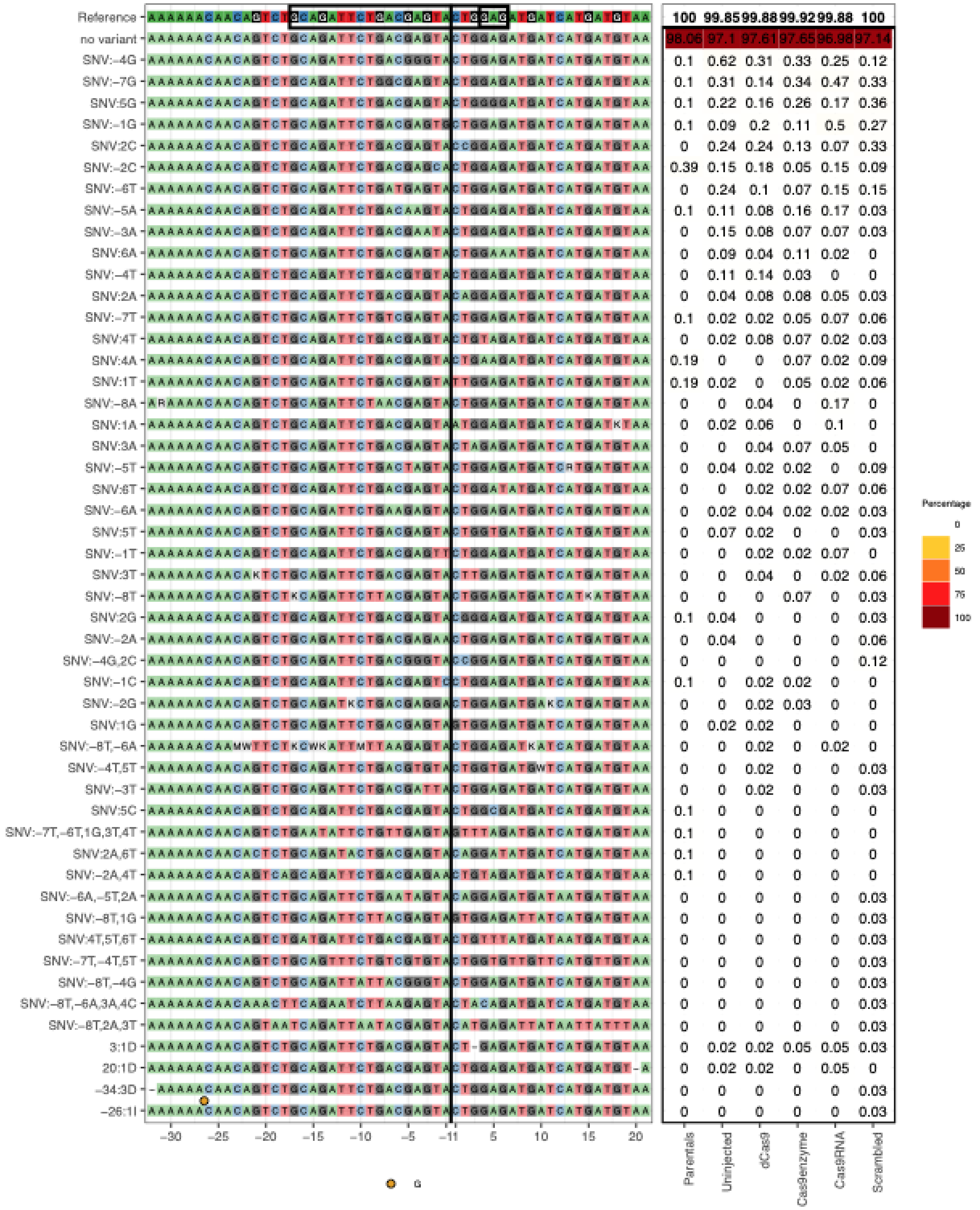
Reference
no variant
Parentals
Uninjected
dCas9
Cas9enzyme
Cas9RNA
Scrambled
Percentage
0
25
50
75
100
G

*il15*:

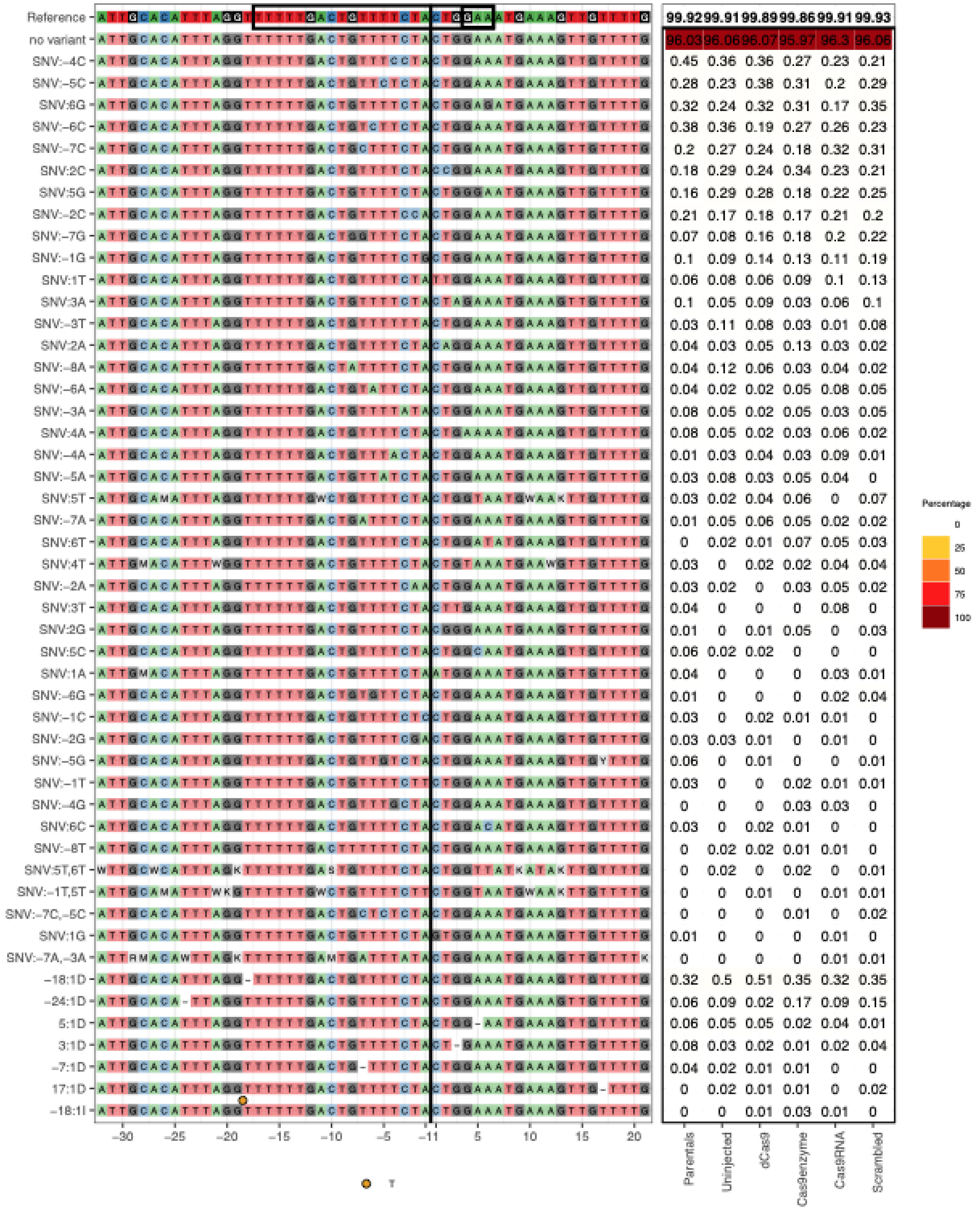
Reference 99.92 99.91 99.89 99.86 99.91 99.93
no variant 96.03 96.06 96.07 95.97 96.3 96.06
SNV:-4C 0.45 0.36 0.36 0.27 0.23 0.21
SNV:-5C 0.28 0.23 0.38 0.31 0.2 0.29
SNV:6G 0.32 0.24 0.32 0.31 0.17 0.35
SNV:-6C 0.38 0.36 0.19 0.27 0.26 0.23
SNV:-7C 0.2 0.27 0.24 0.18 0.32 0.31
SNV:2C 0.18 0.29 0.24 0.34 0.23 0.21
SNV:5G 0.16 0.29 0.28 0.18 0.22 0.25
SNV:-2C 0.21 0.17 0.18 0.17 0.21 0.2
SNV:-7G 0.07 0.08 0.16 0.18 0.2 0.22
SNV:-1G 0.1 0.09 0.14 0.13 0.11 0.19
SNV:1T 0.06 0.08 0.06 0.09 0.1 0.13
SNV:3A 0.1 0.05 0.09 0.03 0.06 0.1
SNV:-3T 0.03 0.11 0.08 0.03 0.01 0.08
SNV:2A 0.04 0.03 0.05 0.13 0.03 0.02
SNV:-8A 0.04 0.12 0.06 0.03 0.04 0.02
SNV:-6A 0.04 0.02 0.02 0.05 0.08 0.05
SNV:-3A 0.08 0.05 0.02 0.05 0.03 0.05
SNV:4A 0.08 0.05 0.02 0.03 0.06 0.02
SNV:-4A 0.01 0.03 0.04 0.03 0.09 0.01
SNV:-5A 0.03 0.08 0.03 0.05 0.04 0
SNV:5T 0.03 0.02 0.04 0.06 0 0.07
SNV:-7A 0.01 0.05 0.06 0.05 0.02 0.02
SNV:6T 0 0.02 0.01 0.07 0.05 0.03
SNV:4T 0.03 0 0.02 0.02 0.04 0.04
SNV:-2A 0.03 0.02 0 0.03 0.05 0.02
SNV:3T 0.04 0 0 0 0.08 0
SNV:2G 0.01 0 0.01 0.05 0 0.03
SNV:5C 0.06 0.02 0.02 0 0 0
SNV:1A 0.04 0 0 0 0.03 0.01
SNV:-6G 0.01 0 0 0 0.02 0.04
SNV:-1C 0.03 0 0.02 0.01 0.01 0
SNV:-2G 0.03 0.03 0.01 0 0.01 0
SNV:-5G 0.06 0 0.01 0 0 0.01
SNV:-1T 0.03 0 0 0.02 0.01 0.01
SNV:-4G 0 0 0 0.03 0.03 0
SNV:6C 0.03 0 0.02 0.01 0 0
SNV:-8T 0 0.02 0.02 0.01 0.01 0
SNV:5T,6T 0 0.02 0 0.02 0 0.01
SNV:-1T,5T 0 0 0.01 0 0.01 0.01
SNV:-7C,-5C 0 0 0 0.01 0 0.02
SNV:1G 0.01 0 0 0 0.01 0
SNV:-7A,-3A 0 0 0 0 0.01 0.01
-18:1D 0.32 0.5 0.51 0.35 0.32 0.35
-24:1D 0.06 0.09 0.02 0.17 0.09 0.15
5:1D 0.06 0.05 0.05 0.02 0.04 0.01
3:1D 0.08 0.03 0.02 0.01 0.02 0.04
-7:1D 0.04 0.02 0.01 0.01 0 0
17:1D 0 0.02 0.01 0.01 0 0.02
-18:1I 0 0 0.01 0.03 0.01 0
-30 -25 -20 -15 -10 -5 -1 1 5 10 15 20
Parentals
Uninjected
dCas9
Cas9enzyme
Cas9RNA
Scrambled
Percentage
0
25
50
75
100
T

*knop1*:

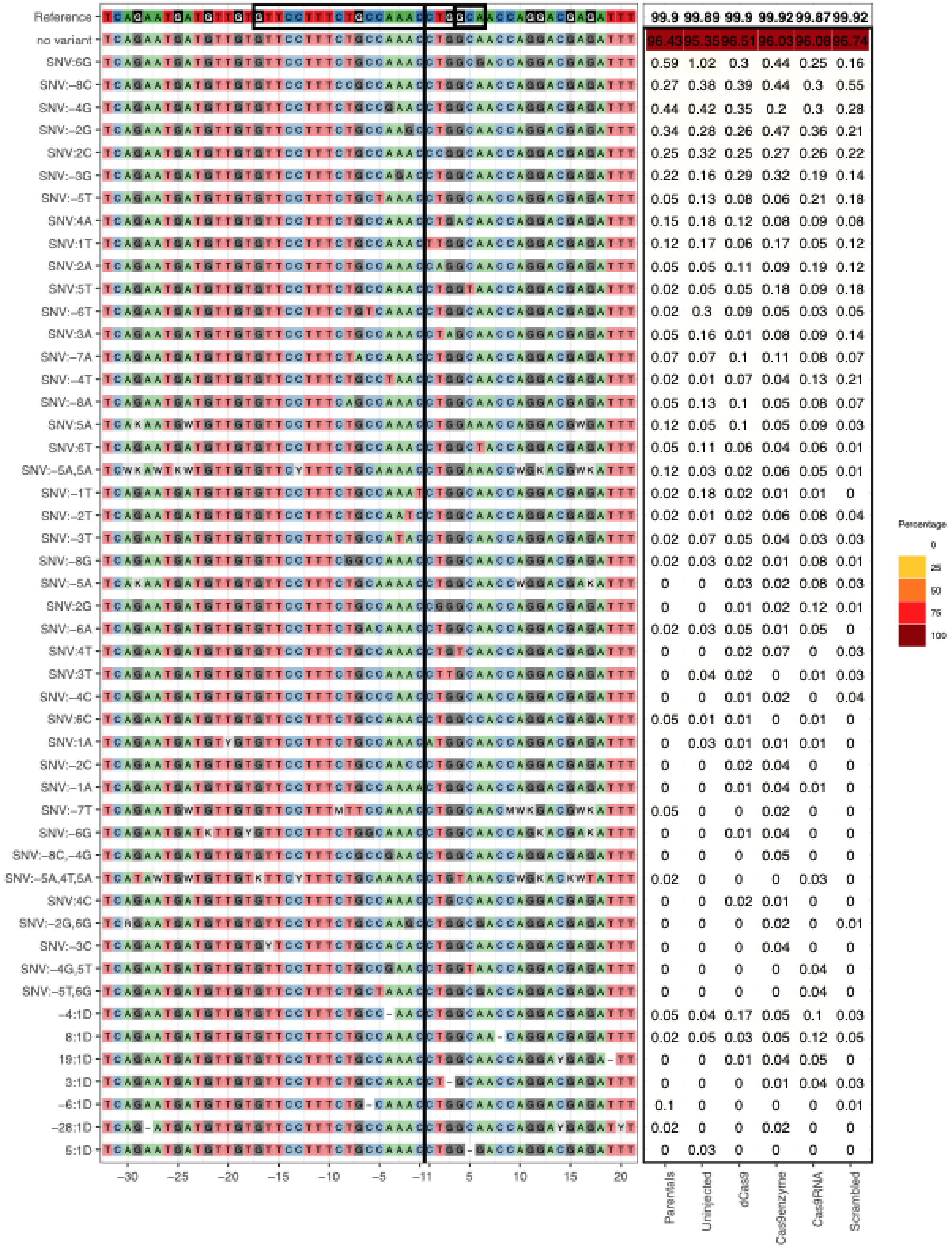
Reference
no variant
99.9 99.89 99.9 99.92 99.87 99.92
Parentals
Uninjected
dCas9
Cas9enzyme
Cas9RNA
Scrambled
Percentage
0
25
50
75
100
−30 −25 −20 −15 −10 −5 −1 1 5 10 15 20

*lamp1a*:

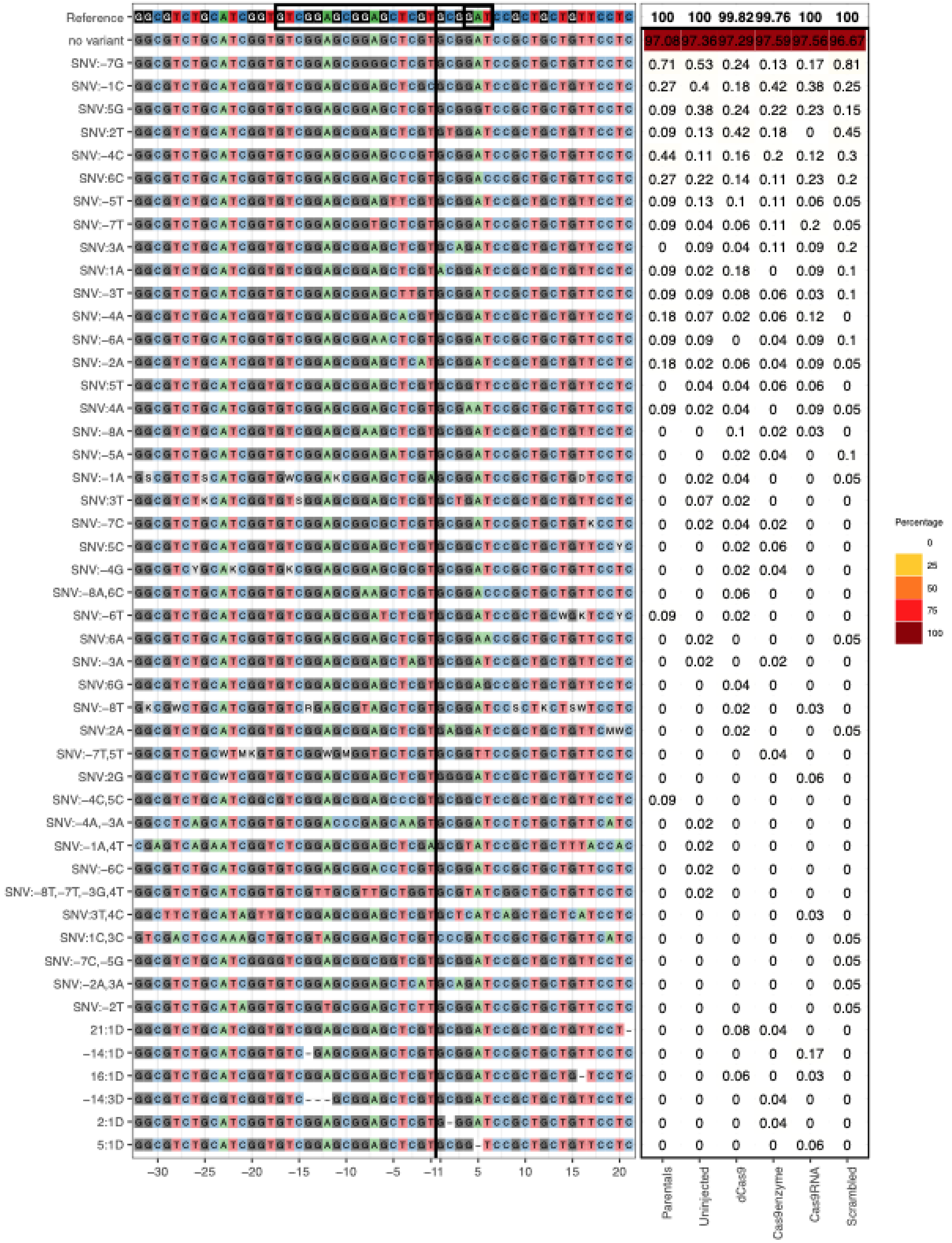

Reference 100 100 99.82 99.76 100 100
no variant 97.08 97.36 97.29 97.59 97.56 96.67
SNV:-7G 0.71 0.53 0.24 0.13 0.17 0.81
SNV:-1C 0.27 0.4 0.18 0.42 0.38 0.25
SNV:5G 0.09 0.38 0.24 0.22 0.23 0.15
SNV:2T 0.09 0.13 0.42 0.18 0 0.45
SNV:-4C 0.44 0.11 0.16 0.2 0.12 0.3
SNV:6C 0.27 0.22 0.14 0.11 0.23 0.2
SNV:-5T 0.09 0.13 0.1 0.11 0.06 0.05
SNV:-7T 0.09 0.04 0.06 0.11 0.2 0.05
SNV:3A 0 0.09 0.04 0.11 0.09 0.2
SNV:1A 0.09 0.02 0.18 0 0.09 0.1
SNV:-3T 0.09 0.09 0.08 0.06 0.03 0.1
SNV:-4A 0.18 0.07 0.02 0.06 0.12 0
SNV:-6A 0.09 0.09 0 0.04 0.09 0.1
SNV:-2A 0.18 0.02 0.06 0.04 0.09 0.05
SNV:5T 0 0.04 0.04 0.06 0.06 0
SNV:4A 0.09 0.02 0.04 0 0.09 0.05
SNV:-8A 0 0 0.1 0.02 0.03 0
SNV:-5A 0 0 0.02 0.04 0 0.1
SNV:-1A 0 0.02 0.04 0 0 0.05
SNV:3T 0 0.07 0.02 0 0 0
SNV:-7C 0 0.02 0.04 0.02 0 0
SNV:5C 0 0 0.02 0.06 0 0
SNV:-4G 0 0 0.02 0.04 0 0
SNV:-8A,6C 0 0 0.06 0 0 0
SNV:-6T 0.09 0 0.02 0 0 0
SNV:6A 0 0.02 0 0 0 0.05
SNV:-3A 0 0.02 0 0.02 0 0
SNV:6G 0 0 0.04 0 0 0
SNV:-8T 0 0 0.02 0 0.03 0
SNV:2A 0 0 0.02 0 0 0.05
SNV:-7T,5T 0 0 0 0.04 0 0
SNV:2G 0 0 0 0 0.06 0
SNV:-4C,5C 0.09 0 0 0 0 0
SNV:-4A,-3A 0 0.02 0 0 0 0
SNV:-1A,4T 0 0.02 0 0 0 0
SNV:-6C 0 0.02 0 0 0 0
SNV:-8T,-7T,-3G,4T 0 0.02 0 0 0 0
SNV:3T,4C 0 0 0 0 0.03 0
SNV:1C,3C 0 0 0 0 0 0.05
SNV:-7C,-5G 0 0 0 0 0 0.05
SNV:-2A,3A 0 0 0 0 0 0.05
SNV:-2T 0 0 0 0 0 0.05
21:1D 0 0 0.08 0.04 0 0
-14:1D 0 0 0 0 0.17 0
16:1D 0 0 0.06 0 0.03 0
-14:3D 0 0 0 0.04 0 0
2:1D 0 0 0 0.04 0 0
5:1D 0 0 0 0 0.06 0
-30 -25 -20 -15 -10 -5 -1 1 5 10 15 20
Parentals
Uninjected
dCas9
Cas9enzyme
Cas9RNA
Scrambled
Percentage
0
25
50
75
100

*nansa*:

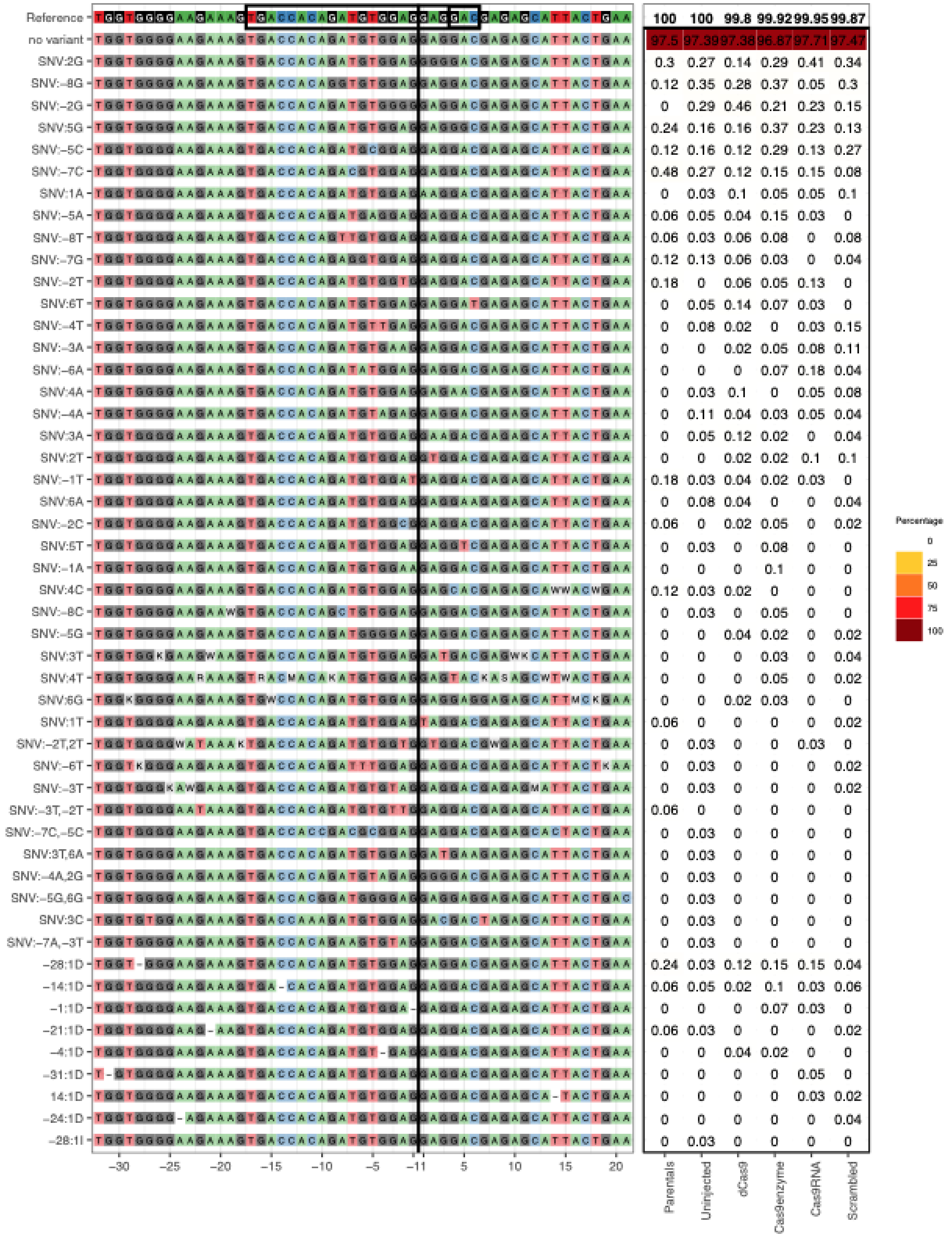
Reference 100 100 99.8 99.92 99.95 99.87
no variant 97.5 97.39 97.38 96.87 97.71 97.47
SNV:2G 0.3 0.27 0.14 0.29 0.41 0.34
SNV:−8G 0.12 0.35 0.28 0.37 0.05 0.3
SNV:−2G 0 0.29 0.46 0.21 0.23 0.15
SNV:5G 0.24 0.16 0.16 0.37 0.23 0.13
SNV:−5C 0.12 0.16 0.12 0.29 0.13 0.27
SNV:−7C 0.48 0.27 0.12 0.15 0.15 0.08
SNV:1A 0 0.03 0.1 0.05 0.05 0.1
SNV:−5A 0.06 0.05 0.04 0.15 0.03 0
SNV:−8T 0.06 0.03 0.06 0.08 0 0.08
SNV:−7G 0.12 0.13 0.06 0.03 0 0.04
SNV:−2T 0.18 0 0.06 0.05 0.13 0
SNV:6T 0 0.05 0.14 0.07 0.03 0
SNV:−4T 0 0.08 0.02 0 0.03 0.15
SNV:−3A 0 0 0.02 0.05 0.08 0.11
SNV:−6A 0 0 0 0.07 0.18 0.04
SNV:4A 0 0.03 0.1 0 0.05 0.08
SNV:−4A 0 0.11 0.04 0.03 0.05 0.04
SNV:3A 0 0.05 0.12 0.02 0 0.04
SNV:2T 0 0 0.02 0.02 0.1 0.1
SNV:−1T 0.18 0.03 0.04 0.02 0.03 0
SNV:6A 0 0.08 0.04 0 0 0.04
SNV:−2C 0.06 0 0.02 0.05 0 0.02
SNV:5T 0 0.03 0 0.08 0 0
SNV:−1A 0 0 0 0.1 0 0
SNV:4C 0.12 0.03 0.02 0 0 0
SNV:−8C 0 0.03 0 0.05 0 0
SNV:−5G 0 0 0.04 0.02 0 0.02
SNV:3T 0 0 0 0.03 0 0.04
SNV:4T 0 0 0 0.05 0 0.02
SNV:6G 0 0 0.02 0.03 0 0
SNV:1T 0.06 0 0 0 0 0.02
SNV:−2T,2T 0 0.03 0 0 0.03 0
SNV:−6T 0 0.03 0 0 0 0.02
SNV:−3T 0 0.03 0 0 0 0.02
SNV:−3T,−2T 0.06 0 0 0 0 0
SNV:−7C,−5C 0 0.03 0 0 0 0
SNV:3T,6A 0 0.03 0 0 0 0
SNV:−4A,2G 0 0.03 0 0 0 0
SNV:−5G,6G 0 0.03 0 0 0 0
SNV:3C 0 0.03 0 0 0 0
SNV:−7A,−3T 0 0.03 0 0 0 0
−28:1D 0.24 0.03 0.12 0.15 0.15 0.04
−14:1D 0.06 0.05 0.02 0.1 0.03 0.06
−1:1D 0 0 0 0.07 0.03 0
−21:1D 0.06 0.03 0 0 0 0.02
−4:1D 0 0 0.04 0.02 0 0
−31:1D 0 0 0 0 0.05 0
14:1D 0 0 0 0 0.03 0.02
−24:1D 0 0 0 0 0 0.04
−28:1I 0 0.03 0 0 0 0
−30 −25 −20 −15 −10 −5 −1 1 5 10 15 20
Parentals
Uninjected
dCas9
Cas9enzyme
Cas9RNA
Scrambled
Percentage
0
25
50
75
100

*si:dkey-9i23.15*:

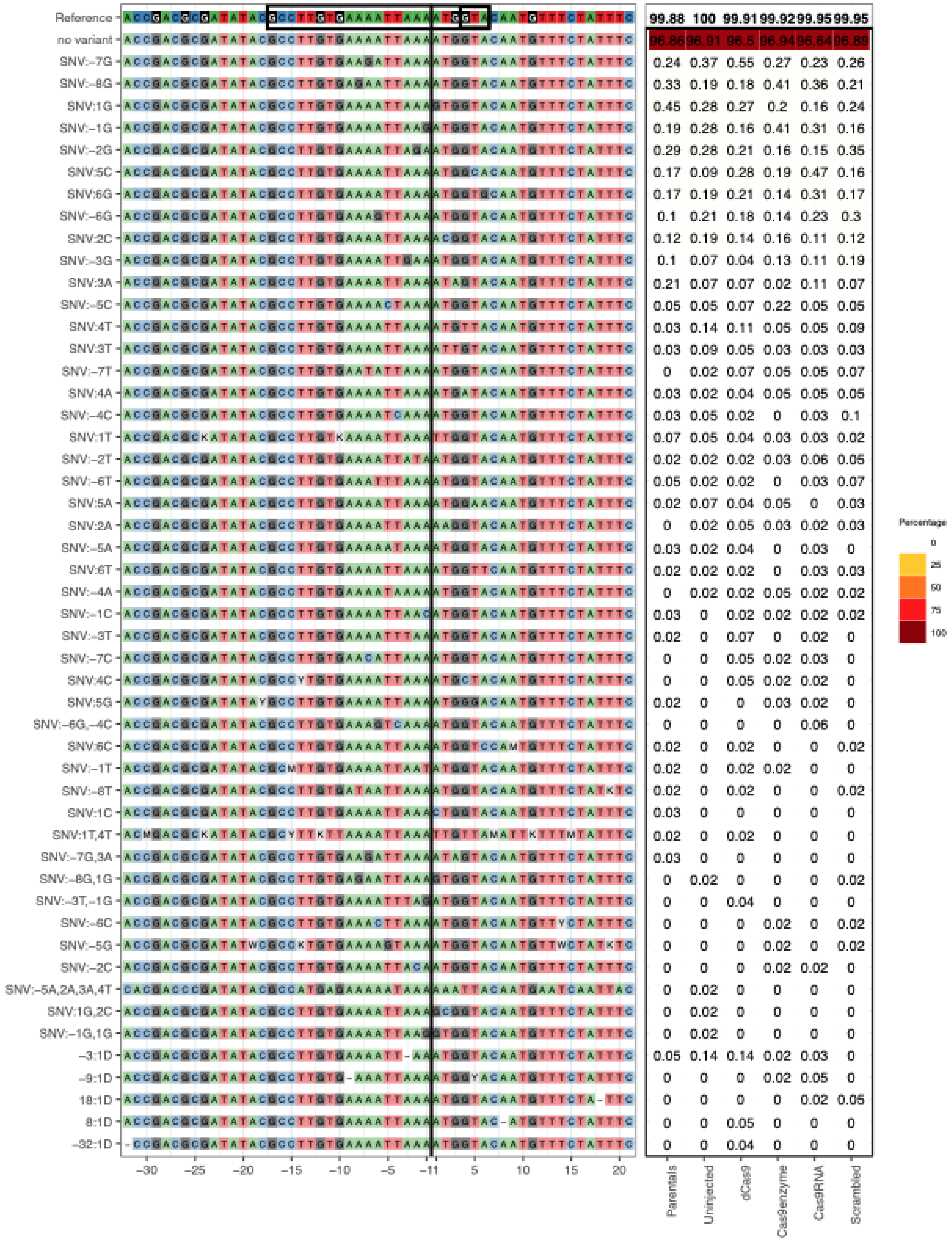

*sich1073220a20:*

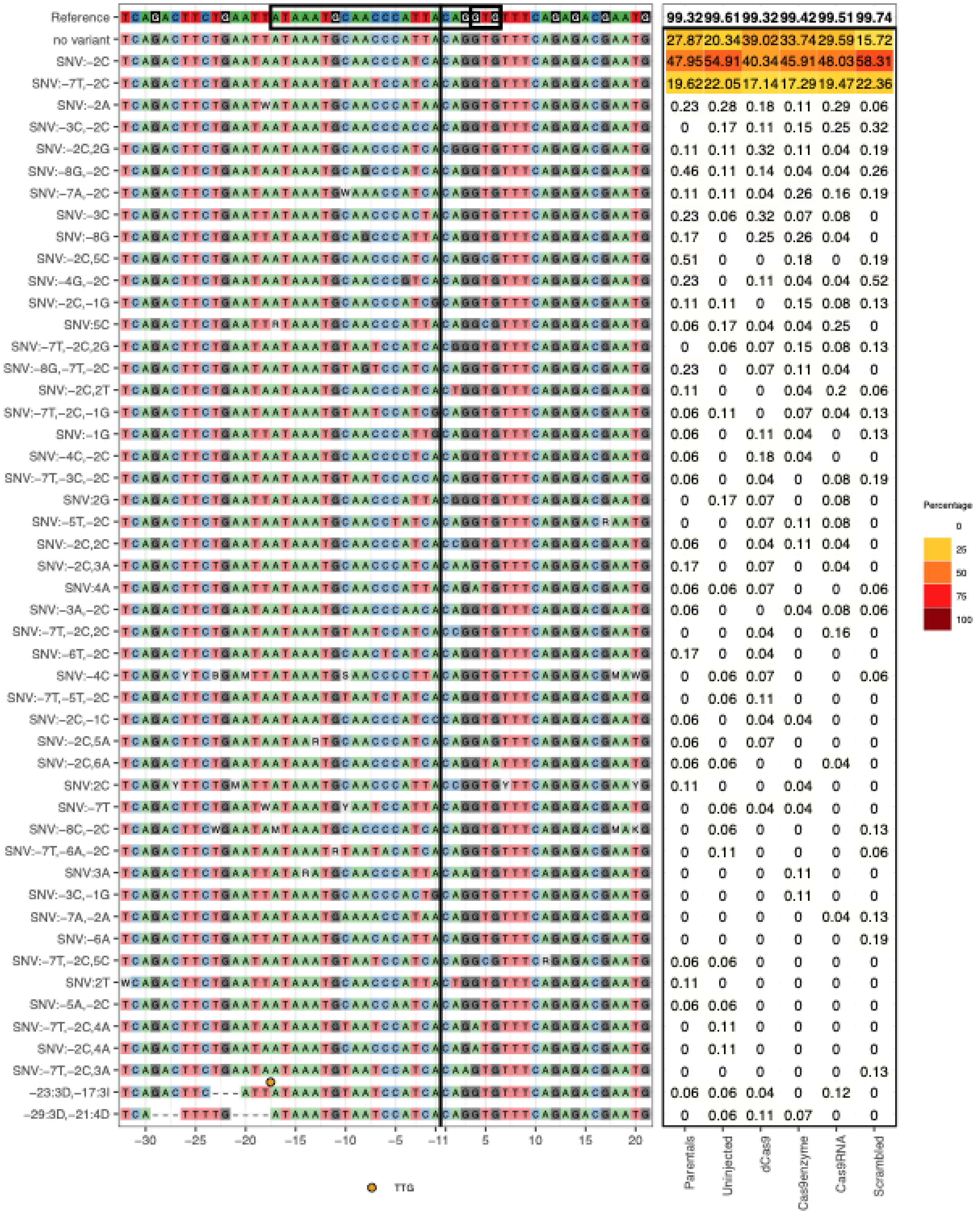

*slc25a4:*

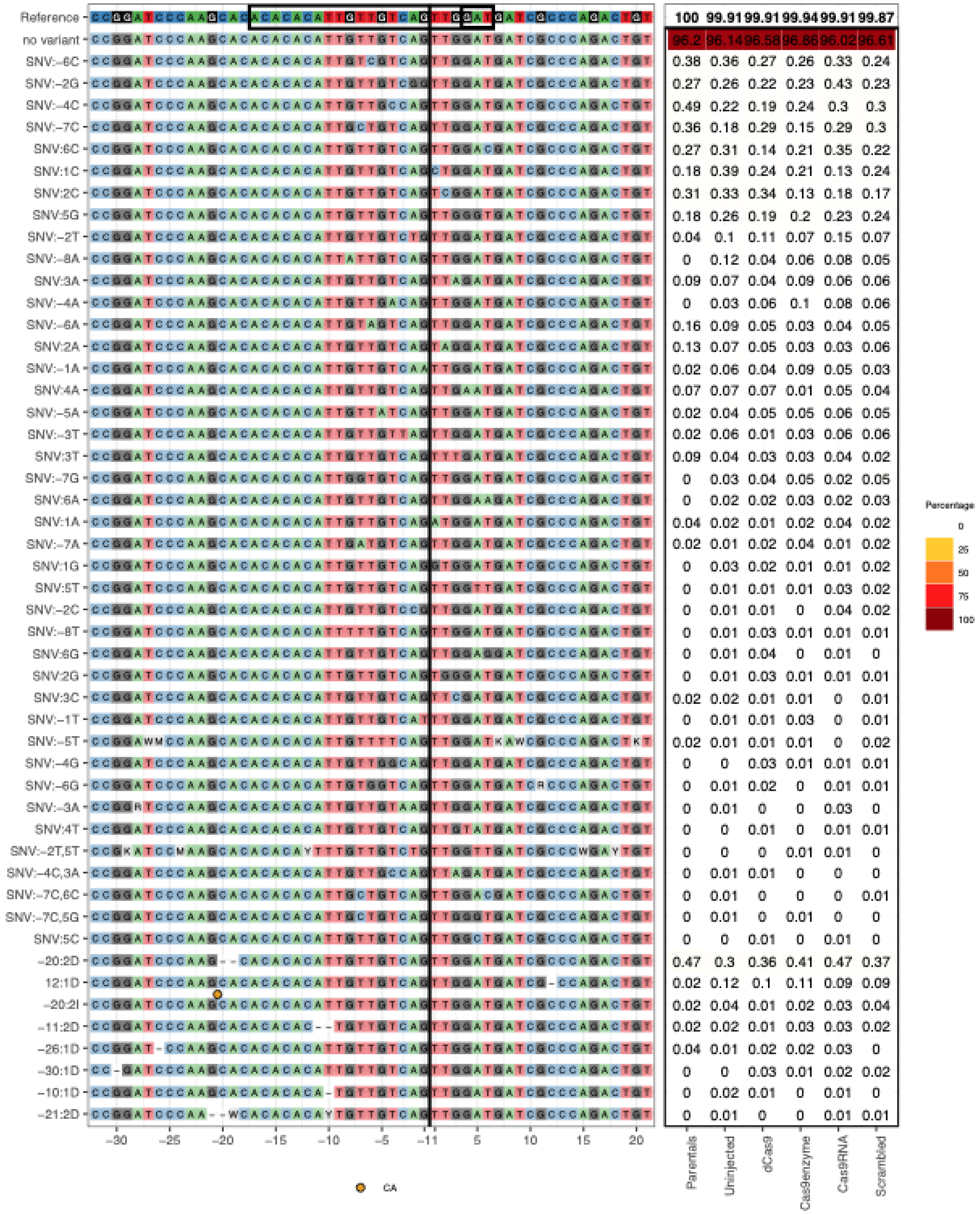
Reference
no variant
SNV:-6C
SNV:-2G
SNV:-4C
SNV:-7C
SNV:6C
SNV:1C
SNV:2C
SNV:5G
SNV:-2T
SNV:-8A
SNV:3A
SNV:-4A
SNV:-6A
SNV:2A
SNV:-1A
SNV:4A
SNV:-5A
SNV:-3T
SNV:3T
SNV:-7G
SNV:6A
SNV:1A
SNV:-7A
SNV:1G
SNV:5T
SNV:-2C
SNV:-8T
SNV:6G
SNV:2G
SNV:3C
SNV:-1T
SNV:-5T
SNV:-4G
SNV:-6G
SNV:-3A
SNV:4T
SNV:-2T,5T
SNV:-4C,3A
SNV:-7C,6C
SNV:-7C,5G
SNV:5C
-20:2D
12:1D
-20:2I
-11:2D
-26:1D
-30:1D
-10:1D
-21:2D
-30 -25 -20 -15 -10 -5 -1 1 5 10 15 20
CA
100 99.91 99.91 99.94 99.91 99.87
96.2 96.14 96.58 96.86 96.02 96.61
0.38 0.36 0.27 0.26 0.33 0.24
0.27 0.26 0.22 0.23 0.43 0.23
0.49 0.22 0.19 0.24 0.3 0.3
0.36 0.18 0.29 0.15 0.29 0.3
0.27 0.31 0.14 0.21 0.35 0.22
0.18 0.39 0.24 0.21 0.13 0.24
0.31 0.33 0.34 0.13 0.18 0.17
0.18 0.26 0.19 0.2 0.23 0.24
0.04 0.1 0.11 0.07 0.15 0.07
0 0.12 0.04 0.06 0.08 0.05
0.09 0.07 0.04 0.09 0.06 0.06
0 0.03 0.06 0.1 0.08 0.06
0.16 0.09 0.05 0.03 0.04 0.05
0.13 0.07 0.05 0.03 0.03 0.06
0.02 0.06 0.04 0.09 0.05 0.03
0.07 0.07 0.07 0.01 0.05 0.04
0.02 0.04 0.05 0.05 0.06 0.05
0.02 0.06 0.01 0.03 0.06 0.06
0.09 0.04 0.03 0.03 0.04 0.02
0 0.03 0.04 0.05 0.02 0.05
0 0.02 0.02 0.03 0.02 0.03
0.04 0.02 0.01 0.02 0.04 0.02
0.02 0.01 0.02 0.04 0.01 0.02
0 0.03 0.02 0.01 0.01 0.02
0 0.01 0.01 0.01 0.03 0.02
0 0.01 0.01 0 0.04 0.02
0 0.01 0.03 0.01 0.01 0.01
0 0.01 0.04 0 0.01 0
0 0.01 0.03 0.01 0.01 0.01
0.02 0.02 0.01 0.01 0 0.01
0 0.01 0.01 0.03 0 0.01
0.02 0.01 0.01 0.01 0 0.02
0 0 0.03 0.01 0.01 0.01
0 0.01 0.02 0 0.01 0.01
0 0.01 0 0 0.03 0
0 0 0.01 0 0.01 0.01
0 0 0 0.01 0.01 0
0 0.01 0.01 0 0 0
0 0.01 0 0 0 0.01
0 0.01 0 0.01 0 0
0 0 0.01 0 0.01 0
0.47 0.3 0.36 0.41 0.47 0.37
0.02 0.12 0.1 0.11 0.09 0.09
0.02 0.04 0.01 0.02 0.03 0.04
0.02 0.02 0.01 0.03 0.03 0.02
0.04 0.01 0.02 0.02 0.03 0
0 0 0.03 0.01 0.02 0.02
0 0.02 0.01 0 0.01 0
0 0.01 0 0 0.01 0.01
Parentals
Uninjected
dCas9
Cas9enzyme
Cas9RNA
Scrambled
Percentage
0
25
50
75
100

*spry2*:

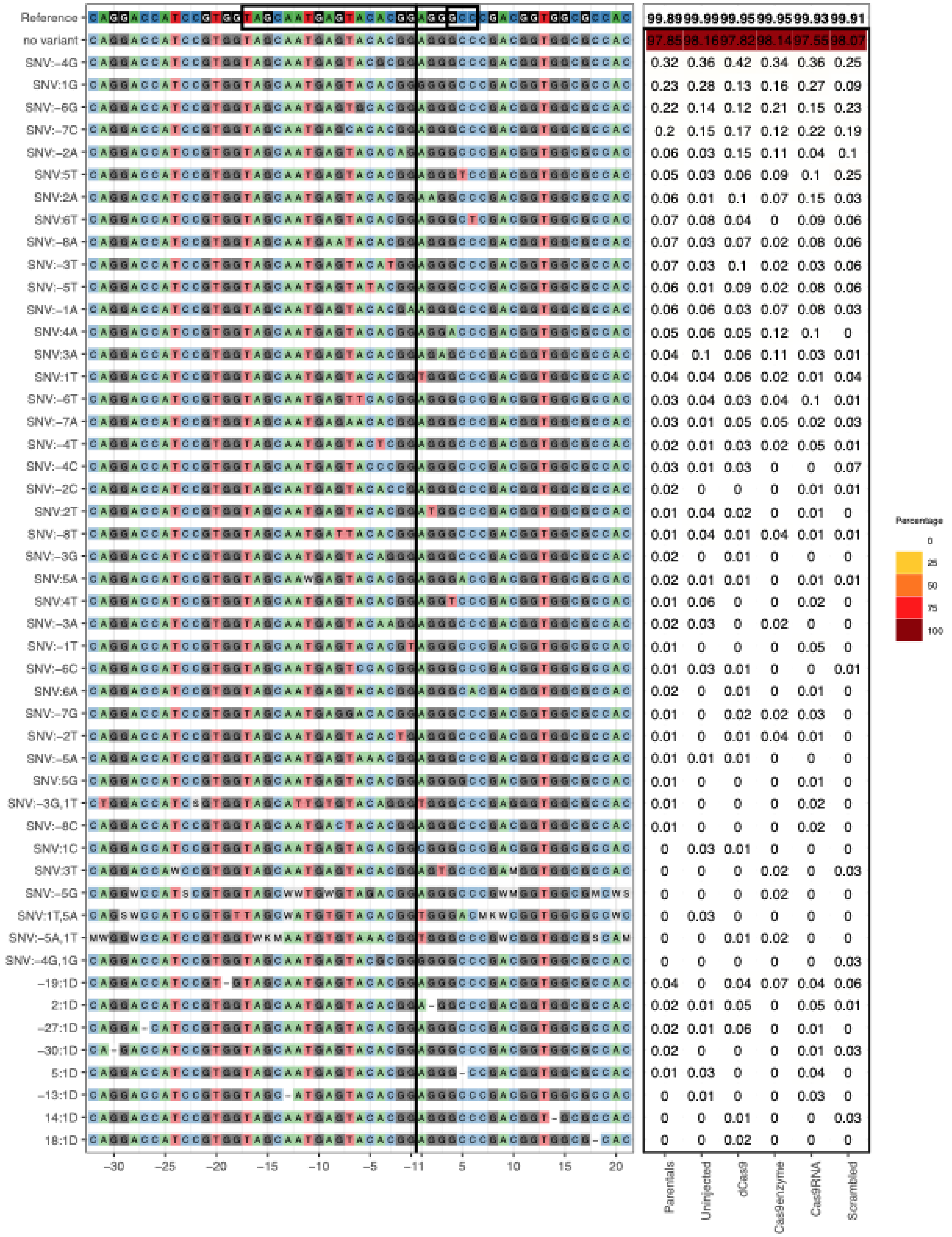

Reference 99.89 99.99 99.95 99.95 99.93 99.91
no variant 97.85 98.16 97.82 98.14 97.55 98.07
SNV:-4G 0.32 0.36 0.42 0.34 0.36 0.25
SNV:1G 0.23 0.28 0.13 0.16 0.27 0.09
SNV:-6G 0.22 0.14 0.12 0.21 0.15 0.23
SNV:-7C 0.2 0.15 0.17 0.12 0.22 0.19
SNV:-2A 0.06 0.03 0.15 0.11 0.04 0.1
SNV:5T 0.05 0.03 0.06 0.09 0.1 0.25
SNV:2A 0.06 0.01 0.1 0.07 0.15 0.03
SNV:6T 0.07 0.08 0.04 0 0.09 0.06
SNV:-8A 0.07 0.03 0.07 0.02 0.08 0.06
SNV:-3T 0.07 0.03 0.1 0.02 0.03 0.06
SNV:-5T 0.06 0.01 0.09 0.02 0.08 0.06
SNV:-1A 0.06 0.06 0.03 0.07 0.08 0.03
SNV:4A 0.05 0.06 0.05 0.12 0.1 0
SNV:3A 0.04 0.1 0.06 0.11 0.03 0.01
SNV:1T 0.04 0.04 0.06 0.02 0.01 0.04
SNV:-6T 0.03 0.04 0.03 0.04 0.1 0.01
SNV:-7A 0.03 0.01 0.05 0.05 0.02 0.03
SNV:-4T 0.02 0.01 0.03 0.02 0.05 0.01
SNV:-4C 0.03 0.01 0.03 0 0 0.07
SNV:-2C 0.02 0 0 0 0.01 0.01
SNV:2T 0.01 0.04 0.02 0 0.01 0
SNV:-8T 0.01 0.04 0.01 0.04 0.01 0.01
SNV:-3G 0.02 0 0.01 0 0 0
SNV:5A 0.02 0.01 0.01 0 0.01 0.01
SNV:4T 0.01 0.06 0 0 0.02 0
SNV:-3A 0.02 0.03 0 0.02 0 0
SNV:-1T 0.01 0 0 0 0.05 0
SNV:-6C 0.01 0.03 0.01 0 0 0.01
SNV:6A 0.02 0 0.01 0 0.01 0
SNV:-7G 0.01 0 0.02 0.02 0.03 0
SNV:-2T 0.01 0 0.01 0.04 0.01 0
SNV:-5A 0.01 0.01 0.01 0 0 0
SNV:5G 0.01 0 0 0 0.01 0
SNV:-3G,1T 0.01 0 0 0 0.02 0
SNV:-8C 0.01 0 0 0 0.02 0
SNV:1C 0 0.03 0.01 0 0 0
SNV:3T 0 0 0 0.02 0 0.03
SNV:-5G 0 0 0 0.02 0 0
SNV:1T,5A 0 0.03 0 0 0 0
SNV:-5A,1T 0 0 0.01 0.02 0 0
SNV:-4G,1G 0 0 0 0 0 0.03
-19:1D 0.04 0 0.04 0.07 0.04 0.06
2:1D 0.02 0.01 0.05 0 0.05 0.01
-27:1D 0.02 0.01 0.06 0 0.01 0
-30:1D 0.02 0 0 0 0.01 0.03
5:1D 0.01 0.03 0 0 0.04 0
-13:1D 0 0.01 0 0 0.03 0
14:1D 0 0 0.01 0 0 0.03
18:1D 0 0 0.02 0 0 0
-30 -25 -20 -15 -10 -5 -1 1 5 10 15 20
Parentals
Uninjected
dCas9
Cas9enzyme
Cas9RNA
Scrambled
Percentage
0
25
50
75
100

*suclg1*:

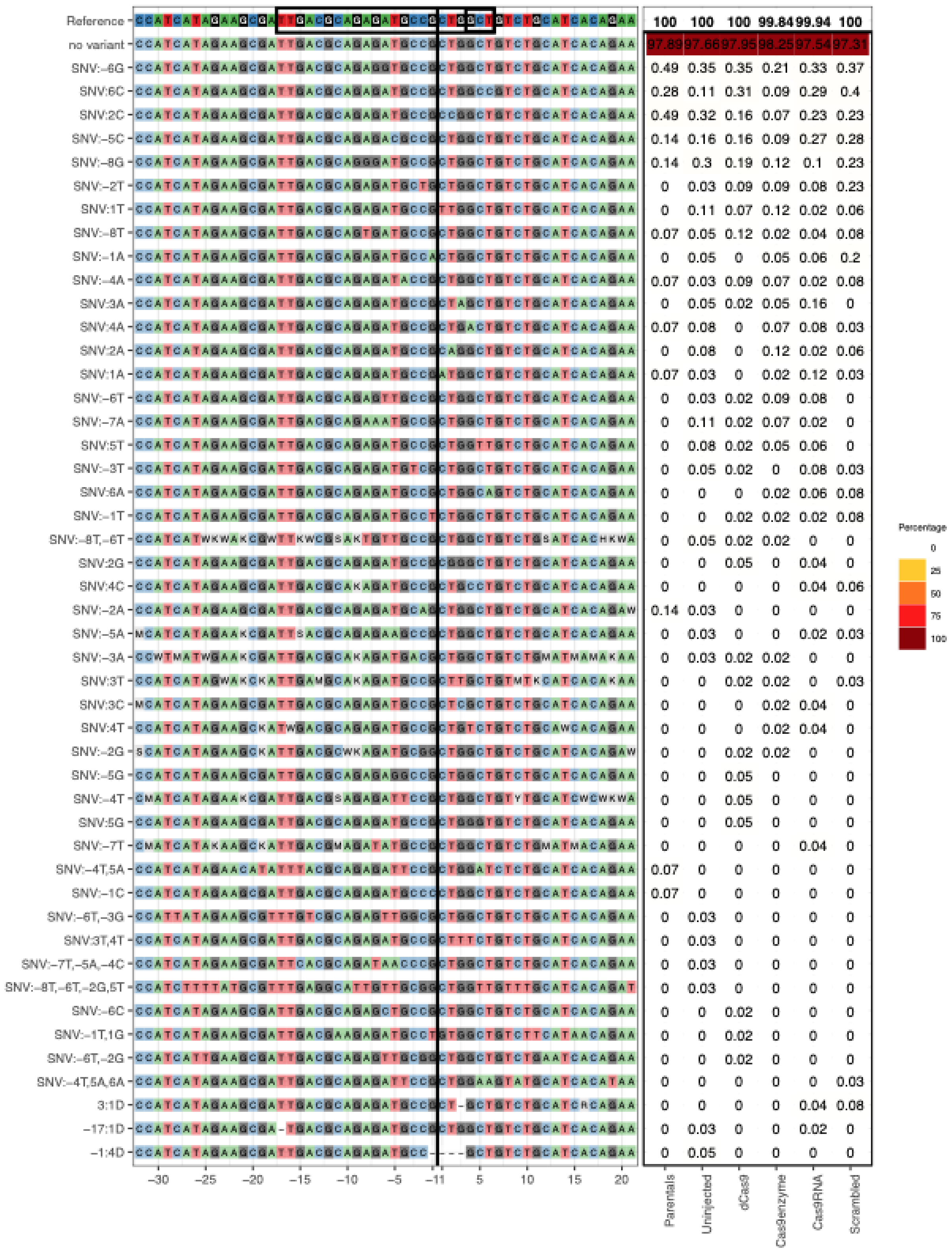
Reference 100 100 100 99.84 99.94 100
no variant 97.89 97.66 97.95 98.25 97.54 97.31
SNV:−6G 0.49 0.35 0.35 0.21 0.33 0.37
SNV:6C 0.28 0.11 0.31 0.09 0.29 0.4
SNV:2C 0.49 0.32 0.16 0.07 0.23 0.23
SNV:−5C 0.14 0.16 0.16 0.09 0.27 0.28
SNV:−8G 0.14 0.3 0.19 0.12 0.1 0.23
SNV:−2T 0 0.03 0.09 0.09 0.08 0.23
SNV:1T 0 0.11 0.07 0.12 0.02 0.06
SNV:−8T 0.07 0.05 0.12 0.02 0.04 0.08
SNV:−1A 0 0.05 0 0.05 0.06 0.2
SNV:−4A 0.07 0.03 0.09 0.07 0.02 0.08
SNV:3A 0 0.05 0.02 0.05 0.16 0
SNV:4A 0.07 0.08 0 0.07 0.08 0.03
SNV:2A 0 0.08 0 0.12 0.02 0.06
SNV:1A 0.07 0.03 0 0.02 0.12 0.03
SNV:−6T 0 0.03 0.02 0.09 0.08 0
SNV:−7A 0 0.11 0.02 0.07 0.02 0
SNV:5T 0 0.08 0.02 0.05 0.06 0
SNV:−3T 0 0.05 0.02 0 0.08 0.03
SNV:6A 0 0 0 0.02 0.06 0.08
SNV:−1T 0 0 0.02 0.02 0.02 0.08
SNV:−8T,−6T 0 0.05 0.02 0.02 0 0
SNV:2G 0 0 0.05 0 0.04 0
SNV:4C 0 0 0 0 0.04 0.06
SNV:−2A 0.14 0.03 0 0 0 0
SNV:−5A 0 0.03 0 0 0.02 0.03
SNV:−3A 0 0.03 0.02 0.02 0 0
SNV:3T 0 0 0.02 0.02 0 0.03
SNV:3C 0 0 0 0.02 0.04 0
SNV:4T 0 0 0 0.02 0.04 0
SNV:−2G 0 0 0.02 0.02 0 0
SNV:−5G 0 0 0.05 0 0 0
SNV:−4T 0 0 0.05 0 0 0
SNV:5G 0 0 0.05 0 0 0
SNV:−7T 0 0 0 0 0.04 0
SNV:−4T,5A 0.07 0 0 0 0 0
SNV:−1C 0.07 0 0 0 0 0
SNV:−6T,−3G 0 0.03 0 0 0 0
SNV:3T,4T 0 0.03 0 0 0 0
SNV:−7T,−5A,−4C 0 0.03 0 0 0 0
SNV:−8T,−6T,−2G,5T 0 0.03 0 0 0 0
SNV:−6C 0 0 0.02 0 0 0
SNV:−1T,1G 0 0 0.02 0 0 0
SNV:−6T,−2G 0 0 0.02 0 0 0
SNV:−4T,5A,6A 0 0 0 0 0 0.03
3:1D 0 0 0 0 0.04 0.08
−17:1D 0 0.03 0 0 0.02 0
−1:4D 0 0.05 0 0 0 0
−30 −25 −20 −15 −10 −5 −1 1 5 10 15 20
Parentals
Uninjected
dCas9
Cas9enzyme
Cas9RNA
Scrambled
Percentage
0
25
50
75
100

*tmem176*:

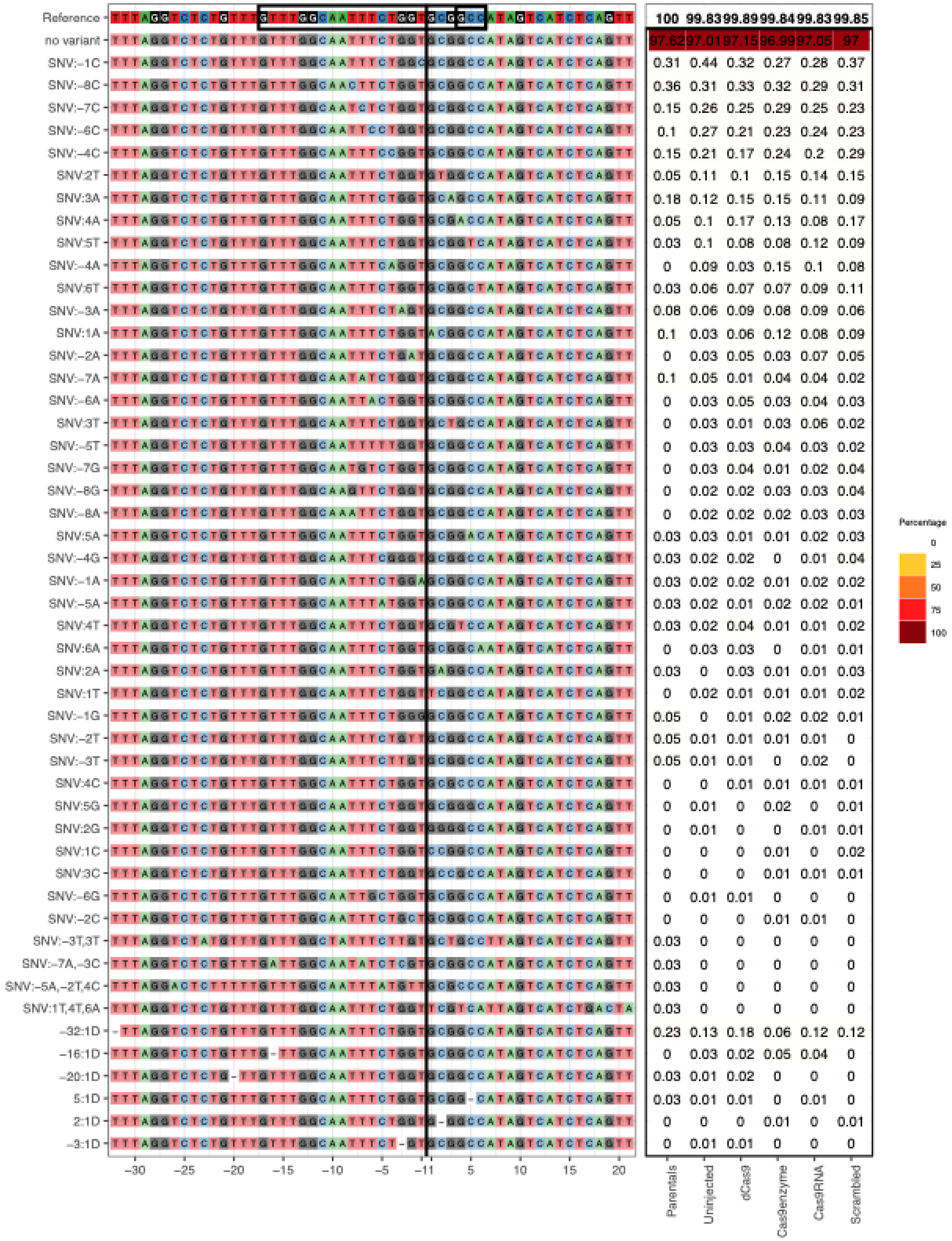
Reference 100 99.83 99.89 99.84 99.83 99.85
no variant 97.62 97.01 97.15 96.99 97.05 97
SNV:–1C 0.31 0.44 0.32 0.27 0.28 0.37
SNV:–8C 0.36 0.31 0.33 0.32 0.29 0.31
SNV:–7C 0.15 0.26 0.25 0.29 0.25 0.23
SNV:–6C 0.1 0.27 0.21 0.23 0.24 0.23
SNV:–4C 0.15 0.21 0.17 0.24 0.2 0.29
SNV:2T 0.05 0.11 0.1 0.15 0.14 0.15
SNV:3A 0.18 0.12 0.15 0.15 0.11 0.09
SNV:4A 0.05 0.1 0.17 0.13 0.08 0.17
SNV:5T 0.03 0.1 0.08 0.08 0.12 0.09
SNV:–4A 0 0.09 0.03 0.15 0.1 0.08
SNV:6T 0.03 0.06 0.07 0.07 0.09 0.11
SNV:–3A 0.08 0.06 0.09 0.08 0.09 0.06
SNV:1A 0.1 0.03 0.06 0.12 0.08 0.09
SNV:–2A 0 0.03 0.05 0.03 0.07 0.05
SNV:–7A 0.1 0.05 0.01 0.04 0.04 0.02
SNV:–6A 0 0.03 0.05 0.03 0.04 0.03
SNV:3T 0 0.03 0.01 0.03 0.06 0.02
SNV:–5T 0 0.03 0.03 0.04 0.03 0.02
SNV:–7G 0 0.03 0.04 0.01 0.02 0.04
SNV:–8G 0 0.02 0.02 0.03 0.03 0.04
SNV:–8A 0 0.02 0.02 0.02 0.03 0.03
SNV:5A 0.03 0.03 0.01 0.01 0.02 0.03
SNV:–4G 0.03 0.02 0.02 0 0.01 0.04
SNV:–1A 0.03 0.02 0.02 0.01 0.02 0.02
SNV:–5A 0.03 0.02 0.01 0.02 0.02 0.01
SNV:4T 0.03 0.02 0.04 0.01 0.01 0.02
SNV:6A 0 0.03 0.03 0 0.01 0.01
SNV:2A 0.03 0 0.03 0.01 0.01 0.03
SNV:1T 0 0.02 0.01 0.01 0.01 0.02
SNV:–1G 0.05 0 0.01 0.02 0.02 0.01
SNV:–2T 0.05 0.01 0.01 0.01 0.01 0
SNV:–3T 0.05 0.01 0.01 0 0.02 0
SNV:4C 0 0 0.01 0.01 0.01 0.01
SNV:5G 0 0.01 0 0.02 0 0.01
SNV:2G 0 0.01 0 0 0.01 0.01
SNV:1C 0 0 0 0.01 0 0.02
SNV:3C 0 0 0 0.01 0.01 0.01
SNV:–6G 0 0.01 0.01 0 0 0
SNV:–2C 0 0 0 0.01 0.01 0
SNV:–3T,3T 0.03 0 0 0 0 0
SNV:–7A,–3C 0.03 0 0 0 0 0
SNV:–5A,–2T,4C 0.03 0 0 0 0 0
SNV:1T,4T,6A 0.03 0 0 0 0 0
–32:1D 0.23 0.13 0.18 0.06 0.12 0.12
–16:1D 0 0.03 0.02 0.05 0.04 0
–20:1D 0.03 0.01 0.02 0 0 0
5:1D 0.03 0.01 0.01 0 0.01 0
2:1D 0 0 0 0.01 0 0.01
–3:1D 0 0.01 0.01 0 0 0
–30 –25 –20 –15 –10 –5 –1 1 5 10 15 20
Parentals
Uninjected
dCas9
Cas9enzyme
Cas9RNA
Scrambled
Percentage
0
25
50
75
100

*vps37c*:

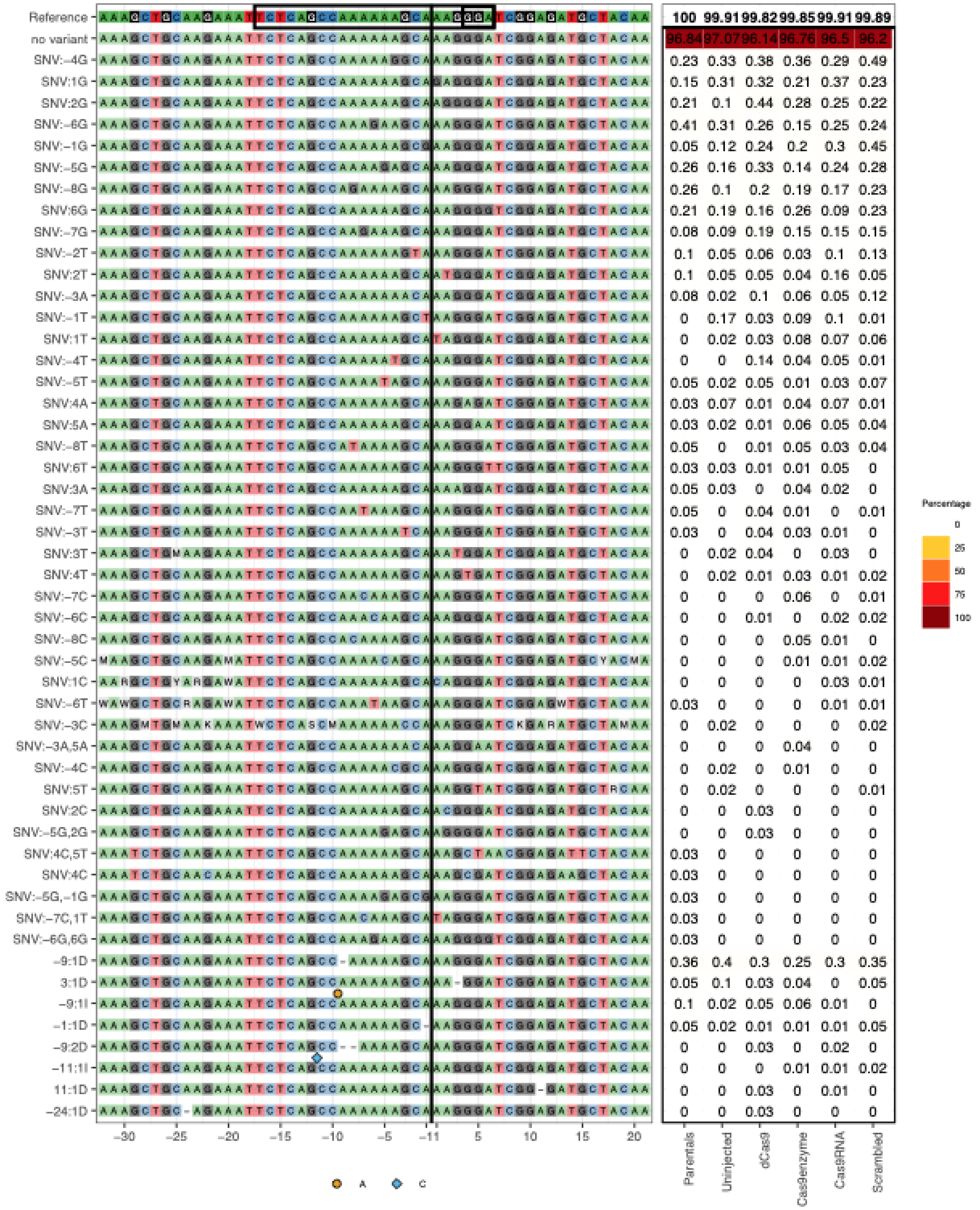
Reference 100 99.91 99.82 99.85 99.91 99.89
no variant 96.84 97.07 96.14 96.76 96.5 96.2
SNV:-4G 0.23 0.33 0.38 0.36 0.29 0.49
SNV:1G 0.15 0.31 0.32 0.21 0.37 0.23
SNV:2G 0.21 0.1 0.44 0.28 0.25 0.22
SNV:-6G 0.41 0.31 0.26 0.15 0.25 0.24
SNV:-1G 0.05 0.12 0.24 0.2 0.3 0.45
SNV:-5G 0.26 0.16 0.33 0.14 0.24 0.28
SNV:-8G 0.26 0.1 0.2 0.19 0.17 0.23
SNV:6G 0.21 0.19 0.16 0.26 0.09 0.23
SNV:-7G 0.08 0.09 0.19 0.15 0.15 0.15
SNV:-2T 0.1 0.05 0.06 0.03 0.1 0.13
SNV:2T 0.1 0.05 0.05 0.04 0.16 0.05
SNV:-3A 0.08 0.02 0.1 0.06 0.05 0.12
SNV:-1T 0 0.17 0.03 0.09 0.1 0.01
SNV:1T 0 0.02 0.03 0.08 0.07 0.06
SNV:-4T 0 0 0.14 0.04 0.05 0.01
SNV:-5T 0.05 0.02 0.05 0.01 0.03 0.07
SNV:4A 0.03 0.07 0.01 0.04 0.07 0.01
SNV:5A 0.03 0.02 0.01 0.06 0.05 0.04
SNV:-8T 0.05 0 0.01 0.05 0.03 0.04
SNV:6T 0.03 0.03 0.01 0.01 0.05 0
SNV:3A 0.05 0.03 0 0.04 0.02 0
SNV:-7T 0.05 0 0.04 0.01 0 0.01
SNV:-3T 0.03 0 0.04 0.03 0.01 0
SNV:3T 0 0.02 0.04 0 0.03 0
SNV:4T 0 0.02 0.01 0.03 0.01 0.02
SNV:-7C 0 0 0 0.06 0 0.01
SNV:-6C 0 0 0.01 0 0.02 0.02
SNV:-8C 0 0 0 0.05 0.01 0
SNV:-5C 0 0 0 0.01 0.01 0.02
SNV:1C 0 0 0 0 0.03 0.01
SNV:-6T 0.03 0 0 0 0.01 0.01
SNV:-3C 0 0.02 0 0 0 0.02
SNV:-3A,5A 0 0 0 0.04 0 0
SNV:-4C 0 0.02 0 0.01 0 0
SNV:5T 0 0.02 0 0 0 0.01
SNV:2C 0 0 0.03 0 0 0
SNV:-5G,2G 0 0 0.03 0 0 0
SNV:4C,5T 0.03 0 0 0 0 0
SNV:4C 0.03 0 0 0 0 0
SNV:-5G,-1G 0.03 0 0 0 0 0
SNV:-7C,1T 0.03 0 0 0 0 0
SNV:-6G,6G 0.03 0 0 0 0 0
-9:1D 0.36 0.4 0.3 0.25 0.3 0.35
3:1D 0.05 0.1 0.03 0.04 0 0.05
-9:1I 0.1 0.02 0.05 0.06 0.01 0
-1:1D 0.05 0.02 0.01 0.01 0.01 0.05
-9:2D 0 0 0.03 0 0.02 0
-11:1I 0 0 0 0.01 0.01 0.02
11:1D 0 0 0.03 0 0.01 0
-24:1D 0 0 0.03 0 0 0
-30 -25 -20 -15 -10 -5 -1 1 5 10 15 20
Parentals
Uninjected
dCas9
Cas9enzyme
Cas9RNA
Scrambled
A
C
Percentage
0
25
50
75
100

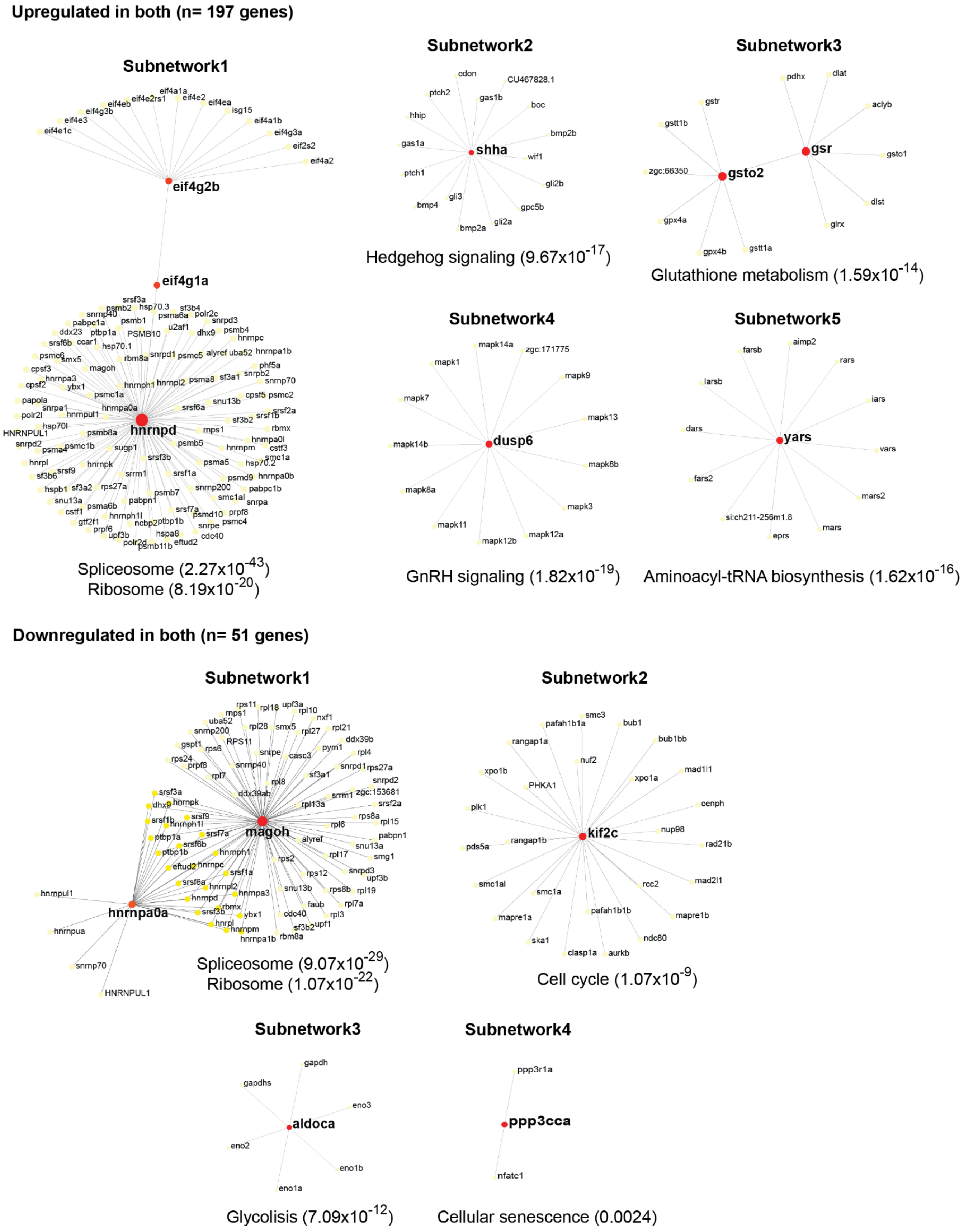


**Supplementary Figure 3**. **Network analyses of the shared differentially expressed genes in Cas9 enzyme and Cas9 mRNA injected larvae.** Each subnetwork includes the top enriched KEGG pathways below with the adjusted *p-values* in parentheses. Network analyses were performed using the Network Analyst online tool (www.networkanalyst.ca).

### Supplementary Tables Legends

**Supplementary Table 1.** Oligonucleotides utilized for all assays performed throughout the project.

**Supplementary Table 2.** On-target efficiency scores for the 50 gRNAs obtained from different *in silico* prediction tools, empirical cleavage scores *in vivo* using Sanger sequencing and sequence deconvolution tools (TIDE and ICE), from the *in vitro* assay CIRCLE-seq (RPMN), and Illumina sequencing using the CrispRVariants software. RPMN: reads per million normalized (see Methods).

**Supplementary Table 3.** Description of the off-targets predicted by CRISPRScan and CIRCLE-seq for the 50 evaluated gRNAs. The percentage of sites intersecting genes from the CRISPRScan tool was obtained from the top 30 predicted off-targets since the tool only provides detailed information from these.

**Supplementary Table 4.** List of the frameshift variants identified in protein-coding genes in samples injected with either Cas9 enzyme or Cas9 mRNA using uninjected batch-siblings as reference (see Methods).

**Supplementary Table 5.** Illumina mosaicism percentages observed in larvae injected with Cas9 (enzyme or mRNA), catalytically dead Cas9 (dCas9), scrambled gRNA, uninjected batch siblings, and a finclip from their crossing parents.

**Supplementary Table 6.** List of differentially expressed genes observed in Cas9-injected samples relative to uninjected batch-siblings and siblings from another batch.

**Supplementary Table 7.** List of gene ontology terms enriched in upregulated genes found in both injection treatments (Cas9 enzyme and Cas9 mRNA).
